# Short 5’ UTRs serve as a marker for viral mRNA translation inhibition by the IFIT2-IFIT3 antiviral complex

**DOI:** 10.1101/2025.02.11.637299

**Authors:** Dustin R. Glasner, Candace Todd, Brian Cook, Agustina D’Urso, Shivani Khosla, Elena Estrada, Jaxon D. Wagner, Mason D. Bartels, Pierce Ford, Jordan Prych, Katie Hatch, Brian A. Yee, Kaori M. Ego, Qishan Liang, Sarah R. Holland, James Brett Case, Kevin D. Corbett, Michael S. Diamond, Gene W. Yeo, Mark A. Herzik, Eric L. Van Nostrand, Matthew D. Daugherty

## Abstract

Recognition of “non-self” nucleic acids, including cytoplasmic dsDNA, dsRNA, or mRNAs lacking proper 5’ cap structures, is critical for the innate immune response to viruses. Here, we demonstrate that short 5’ untranslated regions (UTRs), a characteristic of many viral mRNAs, can also serve as a molecular pattern for innate immune recognition via the interferon-induced proteins IFIT2 and IFIT3. The IFIT2-IFIT3 heterodimer, formed through an intricate domain swap structure resolved by cryo-EM, mediates viral mRNA 5’ end recognition, translation inhibition, and ultimately antiviral activity. Critically, 5’ UTR lengths <50 nucleotides are necessary and sufficient to sensitize an mRNA to translation inhibition by the IFIT2-IFIT3 complex. Accordingly, diverse viruses whose mRNAs contain short 5’ UTRs, such as vesicular stomatitis virus and parainfluenza virus 3, are sensitive to IFIT2-IFIT3-mediated antiviral activity. Our work thus reveals a pattern of antiviral nucleic acid immune recognition that takes advantage of the inherent constraints on viral genome size.

## MAIN

Recognition of foreign RNA is a critical component of immune sensing during viral infection. Interferon (IFN) serves as the first line of defense against viral infection, and its induction is triggered by one of several mechanisms sensing “non-self” RNA or DNA in host cells. IFN signaling induces expression of a number of proteins (e.g., OAS, MDA5, and PKR), which themselves bind to double-stranded RNA (dsRNA) in the cytoplasm to amplify innate immune signaling and activation, promote RNA degradation, or inhibit mRNA translation^1, 2, 3, 4, 5^. Analogously, dsDNA sensing in the cytoplasm by cGAS and other proteins is essential for activation of IFN responses^6, 7, 8^. Other RNAs can be recognized as foreign, including RNAs with an uncapped 5’ tri- or di-phosphate end by RIG-I^1, 2, 3^, mRNAs with high CG-dinucleotide content by zinc finger antiviral protein (ZAP/PARP13)^9^, and endosomal single-stranded RNA (ssRNA) by TLR7 and TLR8^10^. Notably, some of these patterns are also found on “self” nucleic acids, including dsDNA and CG-dinucleotides. Despite this, these nucleic acid sensing proteins comprise a multifaceted barrier to infection by a wide range of viruses.

Among the many RNA-sensing innate immune proteins are the IFIT (IFN-induced proteins with tetratricopeptide repeats) proteins, a family of genes that is highly upregulated during viral infection. Vertebrates encode a varying repertoire of IFITs, with humans encoding five IFIT genes (*IFIT1*, *IFIT1B*, *IFIT2*, *IFIT3*, *IFIT5*), and mice encoding six *(IFIT1*, *IFIT1B*, *IFIT1C*, *IFIT2*, *IFIT3*, *IFIT3B*)^11, 12, 13^. Notably, IFIT1 and IFIT1B proteins recognize “non-self” methylation patterns on the 5’ cap structures of viral mRNAs, leading to inhibition of their translation^13, 14, 15, 16, 17, 18, 19, 20^. In contrast, the role of IFIT2 and IFIT3 in the antiviral response is less clear. Several studies have implicated IFIT2 and IFIT3 in the restriction of viral infection *in vitro* and *in vivo*^11, 21, 22, 23, 24, 25^, and human IFIT3 interacts with and potentiates the effects of human IFIT1^26^. In other studies, IFIT2 has been shown to have a proviral effect on influenza virus replication^27^ and activate apoptosis^28, 29, 30^, whereas IFIT3 may abrogate IFIT2-induced apoptosis^28, 31^. While these studies indicate that IFIT2 and IFIT3 have important roles in the innate immune response, the manner in which IFIT2 and IFIT3 exert direct antiviral effects has remained unclear.

In this study, we demonstrate that IFIT2 and IFIT3 are necessary and sufficient for antiviral activity against vesicular stomatitis virus (VSV) and parainfluenza virus (PIV3). Using cryo-electron microscopy (cryo-EM), virological experiments, and mRNA translation reporter assays, we demonstrate that IFIT2 and IFIT3 form a stable heterodimeric complex that facilitates recognition and translation inhibition of mRNA from several viruses. We determine that the molecular pattern that leads to IFIT2-IFIT3-mediated inhibition is the presence of short (<50 nucleotide (nt)) 5’ untranslated regions (5’ UTRs), which is a feature of mRNAs from many viral families. Our data elucidate a previously undescribed antiviral role for IFIT2 and IFIT3, and reveal that 5’ UTR length is a marker for non-self recognition of viral mRNAs during the innate immune antiviral response.

## RESULTS

### IFIT2 and IFIT3 combine to exert potent antiviral effects *in vitro*

*IFIT2* and *IFIT3* are among the most highly upregulated IFN-stimulated genes (ISGs) following IFN treatment or viral infection^32^. Studies with knockout mice have shown that IFIT2 and/or IFIT3 are required for antiviral activity against several RNA viruses, including VSV^11, 22, 23, 24, 25, 33^. Based on these results, we first determined whether IFIT2 and IFIT3 contribute to the antiviral effects of type I IFN against VSV in human cells. As a positive control, we also evaluated IFIT1, which has been shown to have antiviral activity against VSV^13, 14, 20^. Knockdown of either *Ifit1*, *Ifit2*, or *Ifit3* using gene-specific siRNAs in the A549 human lung epithelial cell line partially rescued viral replication following IFN-α treatment (**Fig. 1a**, **Extended Data** Fig. 1), suggesting that all three IFITs contribute to – but are not individually the entire driver of – the antiviral effect of type I IFN.

**Fig. 1:**
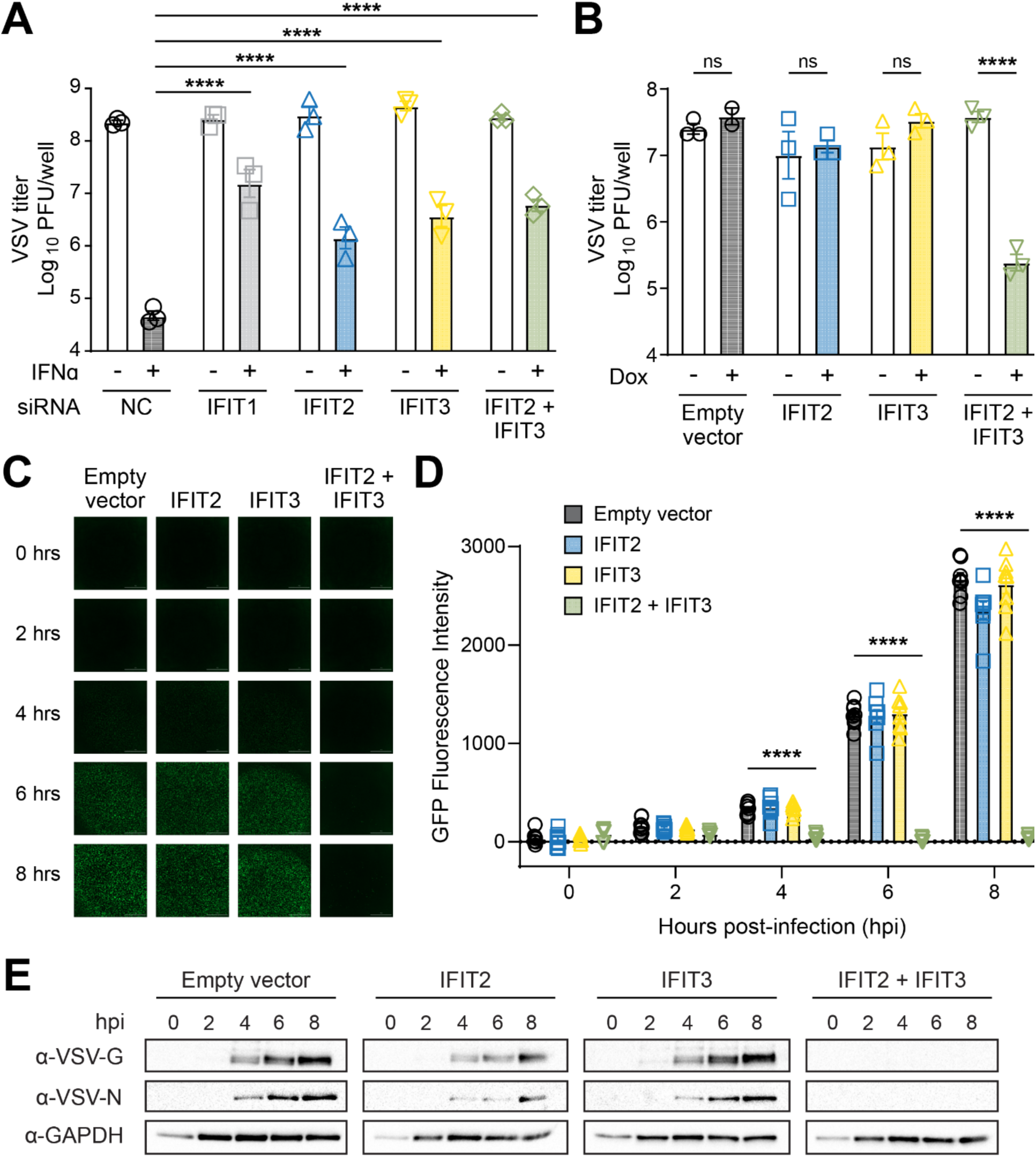
IFIT2 and IFIT3 act together to inhibit viral replication. (**A**) Human A549 cells were treated with a non-targeting control siRNA (NC, black circles) as a negative control, or siRNA targeting IFIT1 (light grey squares), IFIT2 (blue triangles), IFIT3 (yellow triangles), or IFIT2 and IFIT3 in combination (green diamonds). In indicated conditions, cells were induced with IFN-α (1000 U/mL). All cells were infected with VSV-GFP (0.01 MOI), and supernatant was collected at 16 hpi for titration. (**B**) Inducible Flp-In T-REx HEK293 cells expressing no IFIT (black circles), mouse IFIT2 (blue squares), mouse IFIT3 (yellow triangles), or mouse IFIT2 and IFIT3 together (green triangles) were mock treated or treated with doxycycline (500 ng/μL) for 24 h and then infected with VSV-GFP (0.01 MOI). Supernatant was harvested 24 hpi for titration. (**C-E**) Using the same cell lines as in as Figure 1B, each cell line was treated with doxycycline (500 ng/μL) for 24 h and infected with VSV-GFP (3.0 MOI). (**C**) Images were taken at 0, 2, 4, 6, and 8 hpi. (**D**) Images from experiments shown in panel C were processed and quantified for GFP fluorescence intensity. (**E**) Lysates from infected cells were harvested at 0, 2, 4, 6, and 8 hpi and analyzed by western blotting for expression of two viral proteins, VSV-G and VSV-N, and a loading control, GAPDH. All experiments were performed with three or more biological replicates with individual data points shown (**A, B, D**) or representative images shown (**C, E**). Statistical analyses: Data are represented as mean ± SEM. (**A, D**) Ordinary two-way ANOVA with Tukey’s post-test and a single pooled variance. **** = p<0.0001, ns = not significant; (**B**) ordinary two-way ANOVA with Šídák’s post-test and a single pooled variance. **** = p<0.0001.

Human IFIT1, IFIT2, and IFIT3 can interact with and regulate one another^14, 26, 34, 35, 36^, making it difficult to determine from the above experiments whether IFIT2 and IFIT3 have antiviral activity distinct from IFIT1. To establish whether IFIT2 and IFIT3, alone or in combination, are sufficient for antiviral activity in the absence of an IFIT1 interaction, we took advantage of the observation that mouse IFIT2 and IFIT3 do not interact with IFIT1^26, 34^. Because HEK293 cells basally express low levels of IFIT2 and IFIT3^12^ (https://www.proteinatlas.org/ENSG00000119922-IFIT2/cell+line; https://www.proteinatlas.org/ENSG00000119917-IFIT3/cell+line) and do not mount functional inflammatory responses^37^, we generated HEK293 cell lines that express mouse IFIT2 and mouse IFIT3 alone or in combination under the inducible control of doxycycline (DOX) (**Extended Data** Fig. 1). Upon DOX induction, we observed no decrease in VSV titers when mouse IFIT2 or IFIT3 was expressed alone (**Fig. 1b)**. However, when mouse IFIT2 and IFIT3 were co-expressed in HEK293 cells, we observed a >1,000-fold decrease in VSV titers (**Fig. 1b)**. Notably, a significant difference in GFP expression from the viral genome can be observed as early as 4 hours post-infection (**Fig. 1c,d)**. Together with our knockdown data, these results indicate that IFIT2 and IFIT3 are both necessary for an effective IFN-mediated antiviral activity against VSV and sufficient to attenuate VSV replication when expressed in the absence of IFN stimulation.

Several IFITs have been shown to bind RNA and repress mRNA translation^13, 18, 19, 34^. We therefore evaluated the effect of mouse IFIT2 and IFIT3 expression on VSV-encoded protein levels during infection. At 0, 2, 4, 6, and 8 h post-infection, infected cells were harvested and analyzed by western blotting. Expression of IFIT2 and IFIT3 together, but not alone, prevented detectable accumulation of VSV-G and -N proteins compared to control cells (**Fig. 1e)**. These data support a model in which IFIT2 and IFIT3 cooperate during the antiviral response to disrupt an early step of viral infection, resulting in lower viral protein expression.

### Structure of the mouse IFIT2-IFIT3 heterodimer reveals the basis for antiviral complex assembly

Although IFIT2 and IFIT3 were previously shown to form a stable heterodimer^34^, the structural basis of IFIT2-IFIT3 complex formation had not been characterized. To determine the basis for its antiviral activity and inform structure-guided mutations to disrupt function, we recombinantly expressed and purified a 1:1 complex of mouse IFIT2 and IFIT3 and determined its structure to 3.2 Å resolution by cryo-EM (**Supplementary Table 1**, **Extended Data** Fig. 2).

IFIT proteins are composed of tandem α-helical tetratricopeptide repeats (TPR) that can fold into superhelical spiral structures^12, 15, 32, 38^. In the case of monomeric IFIT1 and monomeric IFIT5, the superhelical structure is broken into four domains termed subdomain (SD) I, SD II, Pivot, and SD III^15^, where RNA is bound in a conserved cleft in SD II (α-helices 7-14) and the non-TPR Pivot α-helices (α-helices 15-16)^15^. In our structure, mouse IFIT2 and IFIT3 also adopt all-α-helical structures, with IFIT2 containing 22 α-helices and IFIT3 containing 19 α-helices (**Fig. 2a,c**, **Extended Data** Fig. 3). However, in the IFIT2-IFIT3 heterodimer, the two proteins are oriented parallel to one another and associate through a domain swap involving α-helices 7-9 in SD II. The large buried surface area that results from this domain swap (>4,000 Å^2^ per protomer) provides a molecular explanation for the stability of the IFIT2-IFIT3 complex^34^, and is similar to the domain topology of an earlier crystal structure of an IFIT2 homodimer^38^. Importantly, in RNA-bound IFIT1 and IFIT5, α-helices 7-9 form a clamp around the bound 5’ end of the mRNA, with the rest of SD II and the Pivot helices forming a deep positively-charged cleft that cradles the mRNA (**Fig. 2d)** whereas in IFIT2-IFIT3, α-helices 7-9 are domain-swapped, such that the equivalent deep clefts in this complex include structural elements from both IFIT2 and IFIT3. As a consequence, SD II is shorter in IFIT2 and IFIT3 relative to IFIT1 and IFIT5, and the α-helices C-terminal to SD II (α-helices 10-22 in IFIT2 and α-helices 10-19 in IFIT3) form a single continuous superhelix that comprises SD III (**Fig. 2a-c**, **Extended Data** Fig. 3).

**Fig. 2:**
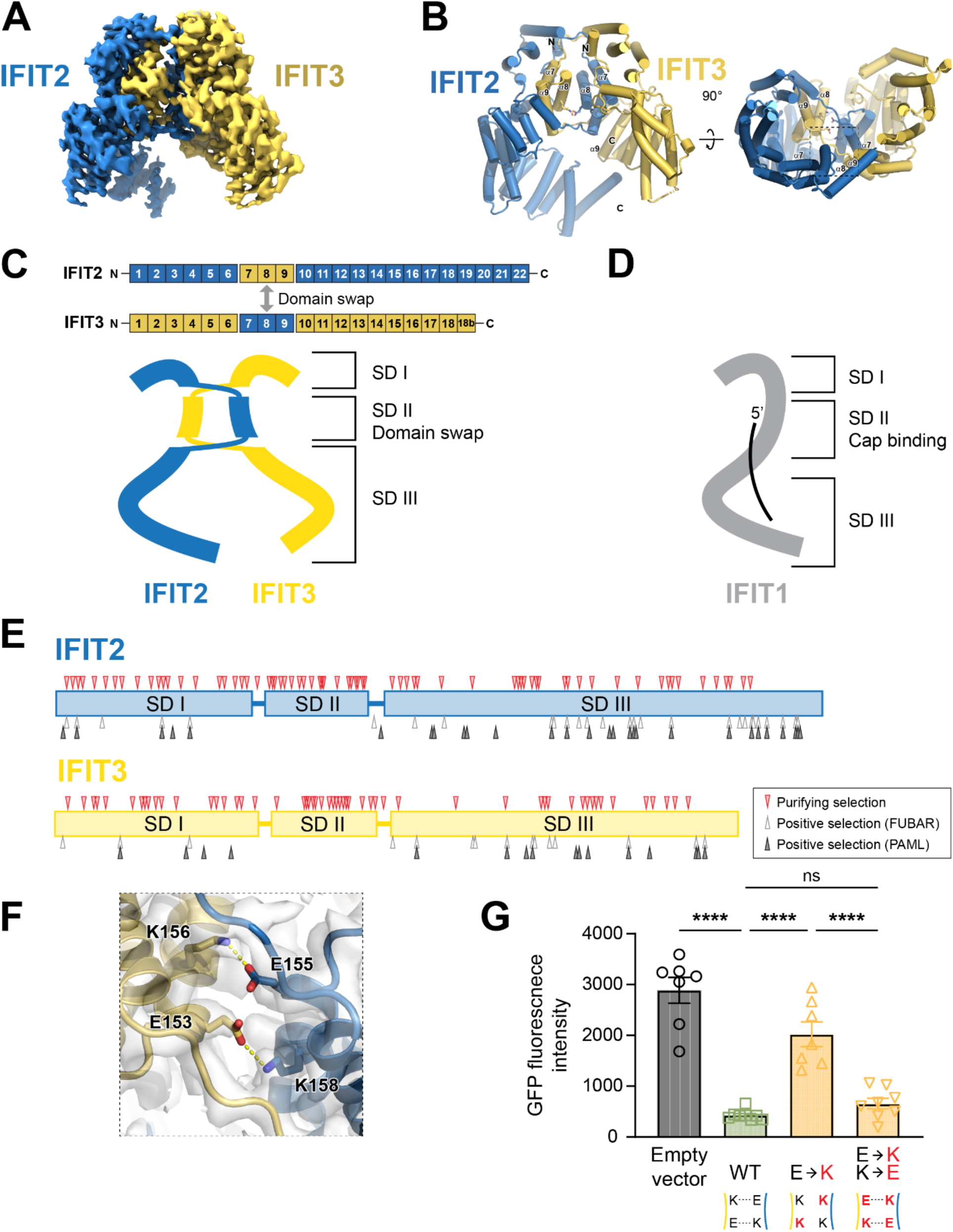
Structure of the IFIT2-IFIT3 heterodimer reveals a conserved interaction surface required for antiviral activity. (**A**) Cryo-EM density map of the mouse IFIT2-IFIT3 heterodimer. (**B**) Structural model of the IFIT2-IFIT3 heterodimer. (**C**) Schematic of the IFIT2-IFIT3 heterodimer, with subdomains (SDs) marked. The SD II domain swap, which forms a continuous superhelix with SD I and SD III, is illustrated. (**D**) Cartoon of the RNA-bound IFIT1 based on PDB 6C6K, highlighting the importance of SD II for 5’ cap binding. (**E**) Evolutionary analyses of IFIT2 and IFIT3 across >20 rodent species, revealing codons evolving under purifying selection (red triangles, from FUBAR analysis) and positive selection (grey and black triangles, from FUBAR and PAML analyses respectively). (**F**) Zoomed-in view of the central helices in SD II that form a reciprocal salt bridge between IFIT2 (blue) and IFIT3 (yellow). Residue numbers are indicated. (**G**) Importance of salt bridge residues for antiviral activity. HEK293T cells were transiently transfected with empty vector, wild-type (WT) IFIT2 and IFIT3, single E->K mutant IFIT2 (E155K) and IFIT3 (E153K), or double mutant E->K/K->E IFIT2 (E155K K158E) and IFIT3 (E153K K156E). Twenty-four hours post-transfection, cells were infected with VSV-GFP (0.05 MOI), and fluorescence images were taken and quantified at 16 hpi. Data (n>5) are represented as mean ± SEM and were analyzed with an ordinary one-way ANOVA with Tukey’s multiple comparisons test. **** = p<0.0001, ns = not significant.

To investigate the importance of the domain-swapped SD II for IFIT2-IFIT3 function, we performed evolutionary analyses on IFIT2 and IFIT3 across species. Consistent with the importance of SD II for function, we observed strong signatures of purifying selection acting on many residues in this region of the protein in both rodents (**Fig. 2e, Supplementary Table 2)** and primates (**Extended Data** Fig. 4, **Supplementary Table 2)**. These evolutionary analyses also revealed strong signatures of positive selection acting on several residues in SD I and especially SD III of both IFIT2 and IFIT3 in both rodents (**Fig. 2e, Supplementary Table 2)** and primates **(Extended Data** Fig. 4**, Supplementary Table 2)**. Such signatures of recurrent positive selection are characteristic of host immunity proteins that are engaged in direct evolutionary arms races with viral antagonists^39, 40, 41, 42^ and similarly strong signatures of positive selection has recently been described for IFIT1^20^. Our evolutionary analyses thus reveal not only the strong conservation of SD II, but also suggest that IFIT2 and IFIT3 are engaged in host-virus arms races as a result of their antiviral activities.

We next evaluated whether mutations at the heterodimer interface in SD II would alter antiviral function. At the core of the SD II domain swap in the IFIT2-IFIT3 heterodimer is a pair of reciprocal salt bridges (**Fig. 2f)**. Wild-type IFIT2 and IFIT3 have glutamic acid residues at amino acid positions 155 and 153, respectively, and lysine residues at positions 158 and 156, respectively. These residues form a pair of E-K salt bridges that we hypothesized are important for complex formation and antiviral activity. To test this idea, we generated single E-to-K mutations in both IFIT2 (E155K) and IFIT3 (E153K). As expected, co-transfection of IFIT2 and IFIT3 single mutant constructs showed less antiviral activity against VSV-GFP than WT IFIT2 and IFIT3, consistent with the predicted K-to-K charge clash at the core of the heterodimeric interface (**Fig. 2g)**. To corroborate this hypothesis, we restored the predicted salt bridges by introducing a second set of K-to-E mutations. Indeed, when we co-transfected these double mutant proteins (IFIT2 E155K/K158E and IFIT3 E153K/K156E), which are predicted to alleviate the charge clash, we restored the antiviral activity of IFIT2-IFIT3 (**Fig. 2g)**. These structural and functional data describe the basis for IFIT2-IFIT3 complex formation and its importance for antiviral activity.

### The IFIT2-IFIT3 antiviral complex binds VSV mRNAs near the start codons

Several IFITs have been described as having RNA binding capabilities ^13, 18, 19, 34^. In the case of IFIT1 and IFIT1B, specific recognition of viral mRNAs, by nature of their “non-self” methylation patterns, is required for their antiviral activity ^13, 14, 18, 19, 20, 34^. Accordingly, we assessed whether IFIT2 or IFIT3, alone or in combination, interacted with specific regions of VSV RNAs by performing eCLIP (enhanced UV-crosslinking and immunoprecipitation) on cells expressing mouse IFIT2 and/or IFIT3 and infected with VSV.

We first performed these experiments in the cells ectopically expressing mouse IFIT2 and IFIT3 alone or in combination and infected with VSV. When IFIT2 and IFIT3 were expressed together, we observed strong immunoprecipitation of RNA sequences toward the 5’ end of the positive (protein-coding) strand of each VSV gene (**Fig. 3a**, **Extended Data** Fig. 5a). The profile of enriched viral RNA was nearly identical for IFIT2 and IFIT3 when co-expressed, whereas we did not observe such enrichment when IFIT2 or IFIT3 was expressed alone. These data further support the model that the IFIT2-IFIT3 heterodimer acts as a single unit, with both IFITs in close proximity to the viral mRNA. Moreover, we observed the strongest enrichment of sequences cross-linking to IFIT2 and IFIT3 in regions that overlap the 5’ UTR and start codons of the VSV mRNA transcripts (**Fig. 3a,b)**. These data suggest that the IFIT2-IFIT3 heterodimer associates with VSV transcripts at sequences proximal to the start codons.

**Fig. 3:**
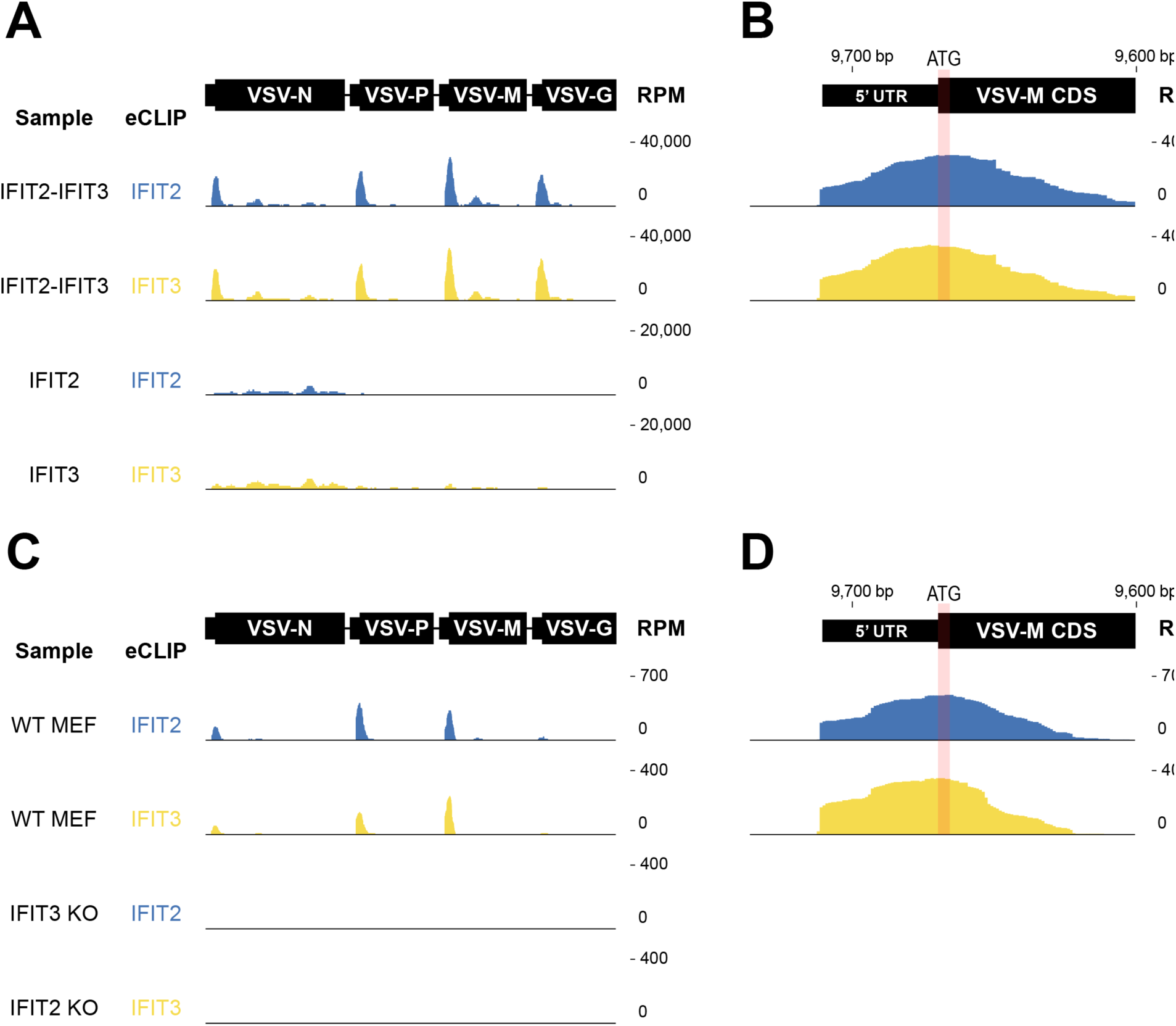
The IFIT2-IFIT3 heterodimer interacts with the 5’ end of VSV mRNAs near the start codons. Browser tracks of VSV reads uniquely mapped to the viral genome as captured by eCLIP of IFIT2 or IFIT3 in (**A-B**) IFIT-expressing Flp-In T-REx HEK293 lines and (**C-D**) wild-type and *Ifit2* or *Ifit3* knockout MEFs. eCLIP data are plotted as reads per million (RPM) mapped across the VSV genome normalized to the total RNA from that region of the viral genome. (**B, D**) Zoomed-in browser tracks for **(B)** 293 and **(D)** MEF lines highlighting peaks of IFIT2-IFIT3 binding proximal to the start codon of the VSV-M gene.

To test whether IFIT2 and IFIT3 associate with the 5’ regions of viral mRNAs during an endogenous IFN response, we performed eCLIP experiments in mouse embryonic fibroblasts (MEFs) in which *Ifit2* or *Ifit3a* and *Ifit3b* had been knocked out (**Fig. 3c,d**, **Extended Data** Fig. 5b). Consistent with our ectopic expression data, we observed that in WT MEFs induced with IFN, IFIT2 and IFIT3 cross-linked with RNA sequences near the start codons of VSV mRNAs. In contrast, this association was lost when either *Ifit2* or *Ifit3a* and *Ifit3b* were knocked out. Although there are differences between our data with ectopically-expressing cells and IFN-induced cells, we observe a similar overall pattern in IFIT2 and IFIT3 association with the 5’ ends of VSV mRNAs and a necessity for both IFIT2 and IFIT3 together during the antiviral response. These data further support that the IFIT2-IFIT3 complex associates with viral mRNAs near their start codons during an innate immune response in infected cells.

### The IFIT2-IFIT3 complex inhibits translation of mRNAs with short viral 5’ UTRs

Both human IFIT1 and mouse IFIT1B inhibit viral mRNA translation by recognizing different methylation patterns on the 5’ cap of viral mRNA^13^. In contrast, our eCLIP data indicate that the IFIT2-IFIT3 complex interacts with mRNA downstream of the cap, at regions near the start codon (**Fig. 3b,d)**. If a specific cap structure is not required for IFIT2-IFIT3 to recognize viral mRNAs, then we reasoned that viral mRNAs transcribed by the host-encoded RNA polymerase II may also be susceptible to IFIT2-IFIT3 mediated repression. To evaluate this idea, we first cloned the 5’ UTR and ORF (open reading frame) of VSV-N, -P, -M, -G, and -L with C-terminal V5 tags into a plasmid that expresses genes from a CMV promoter. We co-transfected these into HEK293T cells in the presence or absence of mouse IFIT2 and IFIT3-expressing plasmids, and then analyzed viral protein expression by western blotting. Despite containing the sequences bound by IFIT2-IFIT3 in our eCLIP experiments, we did not observe changes in viral protein levels in the presence of IFIT2-IFIT3 expression (**Fig. 4a)**.

**Fig. 4:**
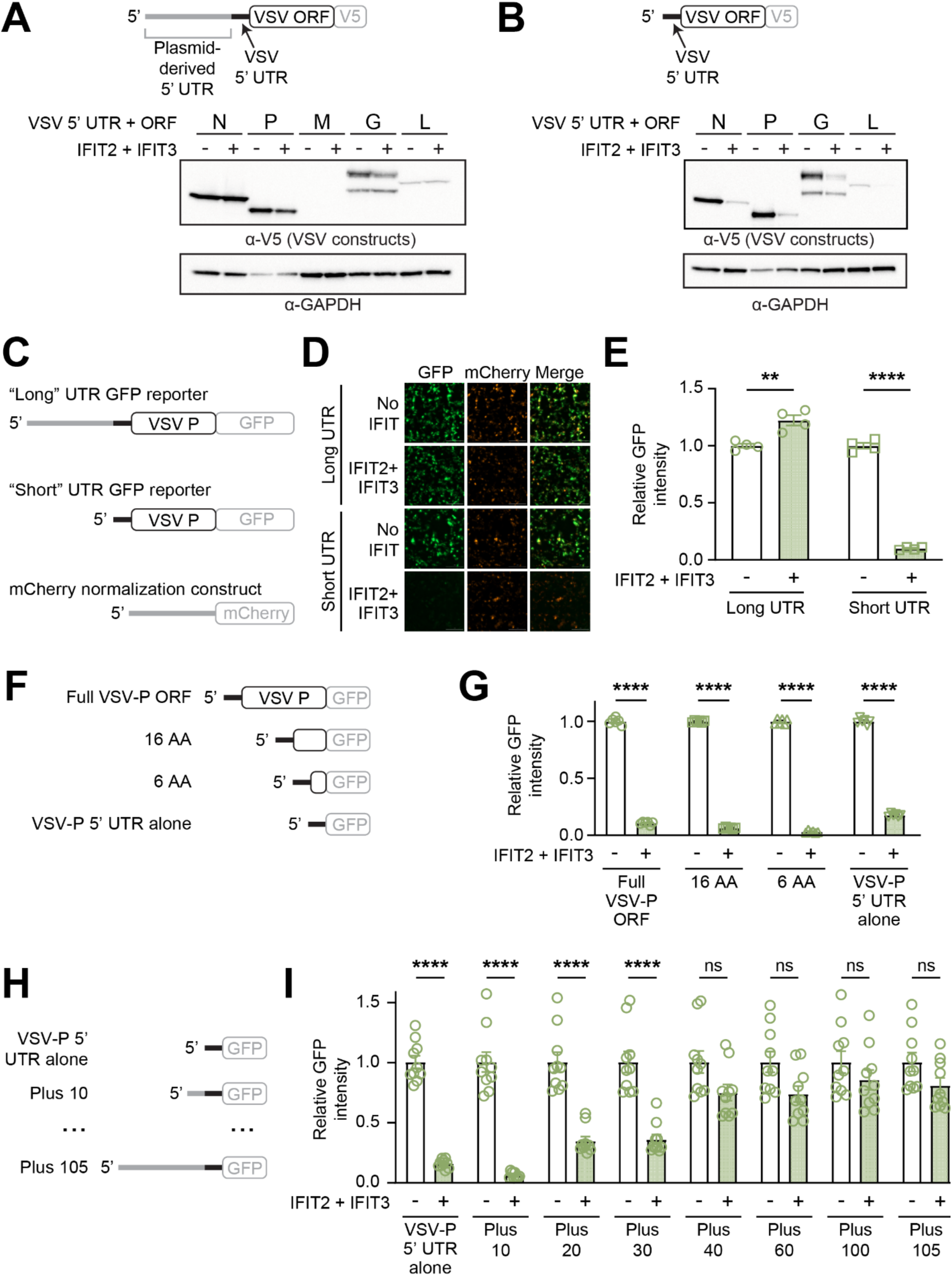
IFIT2-IFIT3 recognizes short viral 5’ UTRs. (**A**) The 5’ UTR and open reading frame (ORF) of each VSV gene was cloned into a mammalian expression plasmid fused to a C-terminal V5 tag (schematic at top). Based on 5’ RACE data **(Extended Data** Fig. 6A), we identified a 108 nt plasmid-derived extension (grey) appended to the 5’ end of the cloned VSV 5’ UTR (black). Each viral gene-expressing construct was co-transfected in the absence or presence of IFIT2-IFIT3 into HEK293T cells. Twenty-four hours post-transfection, cells were harvested and lysates were analyzed by western blotting. (**B**) Following plasmid engineering, only 3 nt of plasmid derived sequence remained at the 5’ end of each VSV UTR and ORF (schematic at top; **Extended Data** Fig. 6B). Transfections and western blotting were performed as in panel A. (**C**) 5’ UTR and ORF schematics for fluorescence reporter constructs shown in panels D and E. (**D**) HEK293T cells were transfected with the indicated GFP reporters in the absence or presence of co-transfected IFIT2-IFIT3. All wells were transfected with the control mCherry normalization construct. Images were taken 24 h post-transfection. (**E**) Experiments were performed as in panel D. For each well (n=4 biological replicates), four images were taken and the ratio of GFP:mCherry signal intensity (see **Extended Data** Fig. 6D) was calculated for each image. All values were normalized to the average of the condition in which IFIT2-IFIT3 was not transfected. (**F**) 5’ UTR and ORF schematics for GFP reporter constructs shown in panel G. AA=amino acids. (**G**) Experiments with the indicated constructs were performed and quantified as in panel E. (**H**) 5’ UTR and ORF schematics for GFP reporter constructs shown in panel I. Plasmid derived sequence from the “long” construct shown in panel C was added back by the indicated number of nt (e.g. Plus 10). (**I**) Experiments with the indicated constructs were performed and quantified as in panel E (n=10 biological replicates). Statistical analyses: Data are represented as mean ± SEM. (**E, G, I**) Ordinary two-way ANOVA with Šídák’s post-test and a single pooled variance. **** = p<0.0001, ns = not significant.

We next considered features of VSV mRNAs near the start codon that could differentiate them from most host mRNAs. One apparent difference is the vast discrepancy in 5’ UTR length between VSV mRNAs, which range from 10 nt to 41 nt, and human mRNAs, which have a median 5’ UTR length of 218 nt^43^. Based on this observation, we hypothesized that 5’ UTR length might serve as a molecular pattern for viral mRNA recognition by IFIT2-IFIT3. We first used 5’ RACE (Rapid Amplification of cDNA Ends) to determine that the transcription start site (TSS) of our plasmid was 105 nt upstream of where we cloned our viral sequences, resulting in a much longer 5’ UTR than natural VSV mRNAs (**Fig. 4a**, **Extended Data** Fig. 6a). Accordingly, we removed this extraneous sequence between the TSS and our viral 5’ UTR, leaving only 3 nt upstream of the viral 5’ UTR that provided a consistent TSS (**Fig. 4b**, **Extended Data** Fig. 6b). With this construct, we now observed a striking reduction in VSV-N, -P, -G, and -L protein expression when IFIT2-IFIT3 was expressed (VSV-M was excluded as it failed to express when transfected, independent of IFIT2-IFIT3) (**Fig. 4b**). These data, which recapitulate the decreased protein expression during viral infection when IFIT2-IFIT3 is expressed (**Fig. 1d**), suggest that 5’ UTR length, not 5’ cap structure, determines the sensitivity of an mRNA to IFIT2-IFIT3-mediated inhibition.

To facilitate more quantitative and higher throughput experiments, we established a fluorescence-based reporter assay. Because VSV-P has the shortest of the VSV 5’ UTRs (10 nt), we cloned eGFP as a C-terminal fusion to VSV-P in our “long” (105 nt plasmid derived UTR followed by 10 nt VSV-P 5’ UTR) and “short” (3 nt followed by 10 nt VSV-P 5’ UTR alone) plasmids (**Fig. 4c**). Consistent with our western blotting data (**Fig. 4a,b**, **Extended Data** Fig. 6c), GFP fluorescence signal was decreased only when it was expressed with a “short” 5’ UTR and IFIT2-IFIT3 was co-expressed (**Fig. 4d**, **Extended Data** Fig. 6d). We quantified a 10-fold decrease in normalized GFP intensity with the short but not the long 5’ UTR construct in cells in which IFIT2-IFIT3 was co-expressed (**Fig. 4e**, **Extended Data** Fig. 6d).

Using this reporter assay, we determined whether any part of the VSV-P ORF contributes to the sensitivity to IFIT2-IFIT3. We truncated the VSV-P ORF to the N-terminal 16 or 6 codons, or removed the entire VSV-P coding sequence, leaving only the 10-nt VSV-P 5’ UTR upstream of the eGFP-V5 ORF (**Fig. 4f)**. These three transcripts were all sensitive to IFIT2-IFIT3 mediated inhibition (**Fig. 4g)**, indicating that only the short viral 5’ UTR is required to sensitize an ORF to repression. Importantly, GFP became insensitive to IFIT2-IFIT3 expression when the 10-nt VSV-P 5’ UTR was lengthened to include the original 105-nt plasmid-derived UTR (**Extended Data** Fig. 6e), and neither IFIT2 nor IFIT3 alone inhibited translation of the VSV-P 5’ UTR alone construct (**Extended Data** Fig. 6f). To confirm that the observed effects on protein expression were due to mRNA translation inhibition rather than mRNA degradation, we demonstrated that IFIT2-IFIT3 expression had no negative effect on levels of the either “short” or “long” mRNA transcripts (**Extended Data** Fig. 6g). Additionally, though the 5’ UTRs for all five VSV genes share a conserved 5’ end sequence (AACAG)^44^, they differ in length and downstream sequence. We therefore confirmed that the 5’ UTRs from VSV-N, -M, -G, and -L are all sensitive in our reporter system as well (**Extended Data** Fig. 6h), further demonstrating that multiple viral 5’ UTRs are susceptible to IFIT2-IFIT3-mediated mRNA translation inhibition.

These data show that 5’ UTRs below a certain length become sensitive to IFIT2-IFIT3 inhibition. To identify the 5’ UTR threshold length determinants of IFIT2-IFIT3-driven translation inhibition, we progressively added length to the sensitive 10 nt VSV-P UTR, increasing it back to the complete plasmid-derived 105 nt (**Fig. 4h)**. Using this approach, we demonstrated that the strongest IFIT2-IFIT3-mediated translation inhibition occurs when the total length of the 5’ UTR is less than 50 nt (**Fig. 4i)**.

### IFIT2-IFIT3 has broad antiviral activity driven by the length of 5’ UTRs

The data above suggests that a 5’ UTR that is <50 nt in length can serve as a molecular pattern that sensitizes an mRNA to an IFIT2-IFIT3 complex. Based on this idea, we hypothesized that mRNA from other viruses with short 5’ UTRs would be sensitive to IFIT2-IFIT3 antiviral activity. We first cloned the 5’ UTRs from rabies virus (RABV) into our reporter construct, as RABV is in the same *Rhabdoviridae* family as VSV. All RABV 5’ UTRs are <50 nt, ranging from 12-30 nt, and we found that all RABV transcripts are sensitive to IFIT2-IFIT3 mediated inhibition (**Fig. 5a)**. These data are consistent with prior work showing that RABV is more pathogenic in mice in which either *Ifit2* or *Ifit3* is knocked out^21, 23^.

**Fig. 5:**
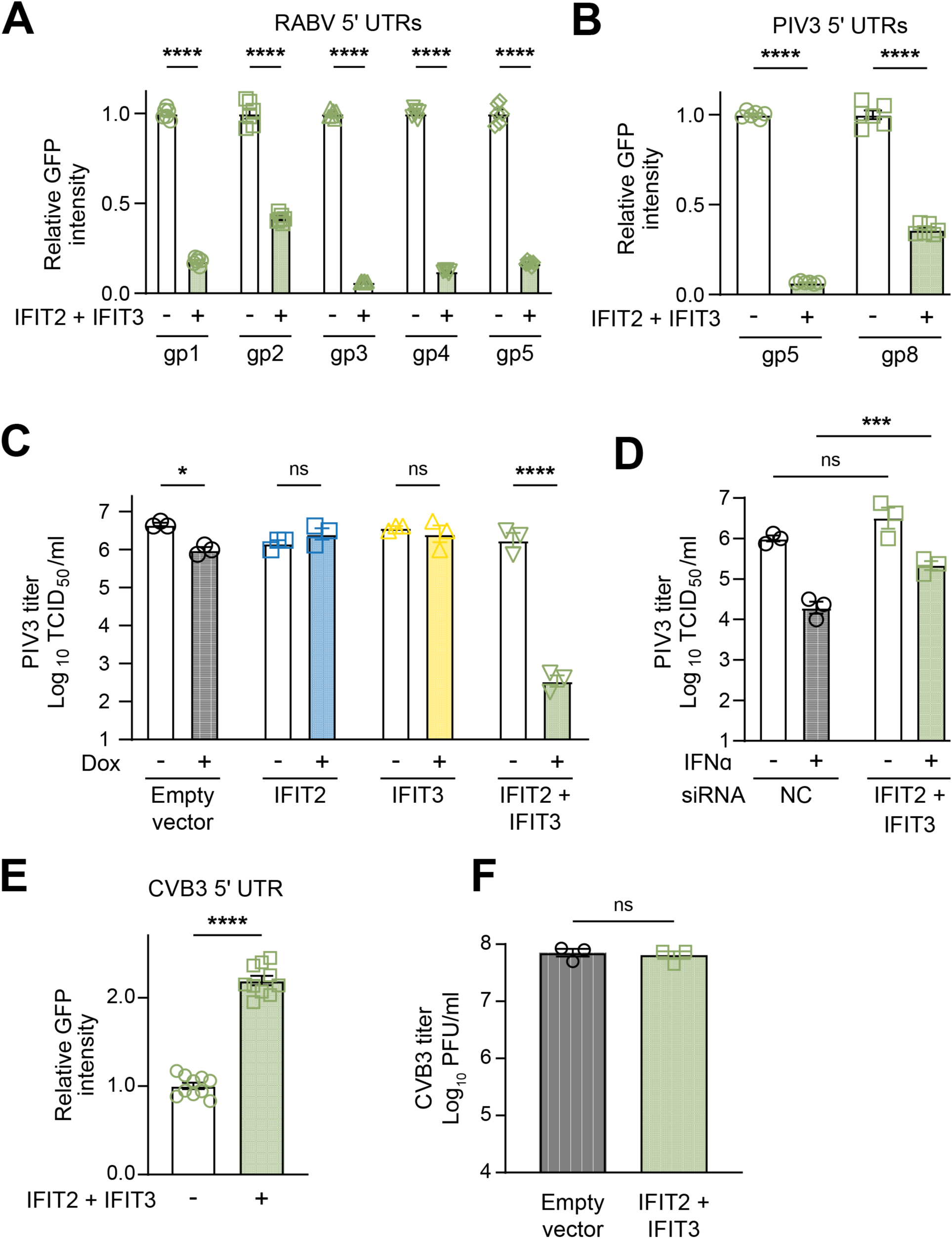
IFIT2-IFIT3 has broad antiviral activity driven by length of 5’ UTRs. (**A-B**) HEK293T cells were transfected with eGFP expression plasmids containing the indicated viral 5’ UTRs in the absence or presence of co-transfected IFIT2-IFIT3. Experiments were performed as in Figure 4E. (**C**) Inducible Flp-In T-REx HEK293 cells expressing no IFIT (black circles), IFIT2 (blue squares), IFIT3 (yellow triangles), or IFIT2 and IFIT3 together (green triangles) were mock-treated or treated with doxycycline (500 ng/μL) for 24 h and then infected with VSV-GFP (0.1 MOI). Supernatant was collected at 48 hpi and quantified by TCID^50^. (**D**) A549 cells were treated with a non-targeting control siRNA (NC, black circles) as a negative control, or siRNA targeting *IFIT2* and *IFIT3* in combination (green diamonds). In indicated conditions, cells were induced with IFN-α (1000 U/mL). All cells were infected with PIV3-GFP (0.1 MOI), and supernatant was collected at 48 hpi for quantification by TCID^50^. (**E**) HEK293T cells were transfected with an eGFP expression plasmid containing the 742 nt CVB3 5’ UTR in the absence or presence of co-transfected IFIT2-IFIT3. (**F**) Inducible Flp-In T-REx HEK293 cells expressing no IFIT (black circles), IFIT2 (blue squares), IFIT3 (yellow triangles), or IFIT2 and IFIT3 together (green triangles) were mock-treated or treated with doxycycline (500 ng/μL) for 24 h and then infected with CVB3 (0.1 MOI). Supernatant was collected at 24 hpi and titered. (**A-F**) All experiments were performed with three or more biological replicates with individual data points shown. Statistical analyses: Data are represented as mean ± SEM. (**A-D**) Ordinary two-way ANOVA with Šídák’s multiple comparisons test and a single pooled variance. * = p<0.05, *** = p<0.001, **** = p<0.0001, ns = not significant; (**E-F**) unpaired parametric t test. **** = p<0.0001, ns = not significant.

To extend our studies, we tested another non-segmented negative-sense RNA virus with several short 5’ UTRs, human parainfluenza virus 3 (PIV3). PIV3 encodes two mRNAs with 5’ UTRs that fall under the 50 nt length threshold (gp5, 32 nt; gp8, 22 nt). When these were tested in our reporter assay, we found that both transcripts were sensitive to IFIT2-IFIT3-mediated inhibition (**Fig. 5b)**. We therefore hypothesized that PIV3 would be sensitive to IFIT2-IFIT3 expression during infection. As with VSV (**Fig. 1b)**, neither IFIT2 nor IFIT3 alone exerted an antiviral effect on PIV3, but their co-expression significantly inhibited infection (**Fig. 5c)**. Also, as with VSV (**Fig. 1a)**, knockdown of *Ifit2* and *Ifit3* expression in human A549 cells blunted the antiviral effects of type I IFN on PIV3 replication (**Fig. 5d)**. Together, our data on VSV, RABV, and PIV3 indicate that IFIT2-IFIT3 co-expression confers inhibitory activity against diverse viruses that encode mRNAs with 5’ UTRs shorter than 50 nt in length.

Finally, we evaluated whether a virus with a long 5’ UTR would be insensitive to the antiviral effects of IFIT2 and IFIT3. We cloned the single 742-nt 5’ UTR of coxsackievirus B3 (CVB3) upstream of eGFP and tested it for sensitivity to IFIT2-IFIT3-mediated translation repression. Indeed, the CVB3 5’ UTR was insensitive to IFIT2-IFIT3 expression, with reporter levels actually increasing (**Fig. 5e)**, suggesting that IFIT2-IFIT3 may be proviral in some contexts as previously shown by Tran et al.^27^. Moreover, as predicted from our reporter system data, ectopic expression of IFIT2 and IFIT3 did not inhibit CVB3 infection (**Fig. 5f)**. These data suggest that viruses with long 5’ UTRs may evade recognition and translation inhibition by the IFIT2-IFIT3 complex.

## DISCUSSION

Recognition of non-self viral mRNA is critical to controlling infection. Host cells must sense and distinguish viral mRNAs to prevent their translation and subsequent viral replication and formation of virions. Here, we show that the ISGs IFIT2 and IFIT3 form a heterodimeric complex capable of exerting a potent antiviral effect against VSV (*Rhabdoviridae*) and PIV3 (*Paramyxoviridae*), and that this activity is driven by recognition and translation inhibition of viral mRNAs containing short 5’ UTRs.

RNA viruses have an average genome length of ∼9 kilobases^45^. With such limited genetic space, many viruses – especially small non-segmented negative strand RNA viruses, encode short 5’ UTRs, presumably to maximize nucleotides available for protein coding sequences. Though RNA viruses have high rates of mutation and can undergo rapid change to genome sequences^46, 47^, the length of viral 5’ UTRs is likely constrained due to constraints on overall genome size^46^, enabling IFIT2 and IFIT3 to have evolved recognition of this feature as a hallmark of RNA virus infection. Indeed, nearly every non-segmented negative-sense RNA virus that infects humans contains at least one mRNA with a 5’ UTR <50 nt, and our data with three of these viruses (PIV3, VSV, and RABV) suggest IFIT2-IFIT3 may be a broad restriction mechanism against them. In comparison, viruses in the henipavirus genus, including Nipah and Hendra viruses, are unique among the *Paramyxoviridae* in not having 5’ UTRs that are <50 nt (ranging from 57 nt to 284 nt). This observation, in addition to our data indicating that the long 5’ UTR of CVB3 and CVB3 infection are insensitive to IFIT2-IFIT3 expression, indicate that evolving longer viral 5’ UTRs to mimic the length of host 5’ UTRs is one potential viral escape mechanism. Combined with studies showing that viruses can evolve RNA secondary structures, host-mimicking RNA methylation machinery, and additional mechanisms to evade IFIT1^16, 20^, these results suggest that IFITs may have had a profound effect on the evolution of the 5’ ends of viruses. Additionally, viruses likely have developed additional strategies to evade translation inhibition by IFIT2-IFIT3. As has been shown with IFIT1^20^, we find that primate and rodent IFIT2 and IFIT3 have evolved under strong recurrent positive selection, especially in SD III. These evolutionary data mirror the strong signatures of positive selection described for other antiviral ISGs that are targeted by viral antagonists^16, 20^, and indicate that some viruses that otherwise might be sensitive to IFIT2-IFIT3-mediated repression likely encode species-specific IFIT antagonists.

The precise mechanism by which the IFIT2-IFIT3 complex selectively inhibits translation of mRNAs with short 5’ UTRs requires further study. Previous models based on ribosome structures and functional data have described a “blind spot” of 40-50 nt at the very 5’ end of human mRNAs, where efficient translation initiation occurs at the first start codon downstream of the blind spot^48^. That viruses with short 5’ UTR-containing mRNAs are able to efficiently translate their proteins during infection suggests that there are exceptions to the blind spot model. As such, viruses that express mRNAs with short 5’ UTRs likely co-opt non-canonical translational pathways. Indeed, VSV translation initiation is strongly dependent on the large ribosomal subunit protein, RPL40 (eL40), whereas bulk cellular translation is not^49^. Likewise, some host mRNAs contain short 5’ UTRs, several of which contain a specific sequence known as a TISU (Translation Initiator of Short 5’ UTR)^50, 51^, which rely on the small ribosomal subunit proteins RPS3 and RPS10^52^, and the initiation factor eIF1 for efficient translation^52, 53^. It is likely that other non-canonical translation mechanisms exist as well. The differential requirements for efficient translation of host mRNAs with short 5’ UTRs leads us to hypothesize that IFIT2 and IFIT3 might selectively inhibit a specialized translation pathway, although further work will be required to understand this mechanism completely.

In summary, our study identifies short mRNA 5’ UTRs as a molecular pattern that at least some mammalian hosts can use to selectively inhibit viral replication. This pattern is present on a wide range of viral mRNAs and is sufficient to sensitize an mRNA to translation inhibition by the IFIT2-IFIT3 antiviral complex. Thus, 5’ UTR length is another point of conflict in the battle between viruses and hosts for mRNA translation control.

## METHODS

### Cell culture and transient transfection

HEK293T cells were obtained from ATCC (catalog # CRL-3216) and grown in complete medium containing DMEM (Gibco, Carlsbad, CA), 10% FBS, 100 U/mL penicillin, and 100 μg/mL streptomycin (Gibco, Carlsbad, CA). For transient transfections, HEK293T cells were seeded the day prior to transfection in a 24-well plate (Genesee, El Cajon, CA) with 500 μL of complete media. Cells were transiently transfected with 500 ng of total DNA and 1.5 μL of TransIT-X2 (Mirus Bio, Madison, WI) following the manufacturer’s protocol.

For generation of inducible cell lines, sequences for mCherry, IFIT2, IFIT3, or IFIT2 and IFIT3 separated by a P2A site were cloned into the Flp-In vector pcDNA5/FRT/TO. Flp-In T-REx HEK293 cells (Invitrogen) maintained in 5 μg/mL of blasticidin were transfected at 70% confluency with mCherry or IFIT constructs and the vector containing the Flp recombinase pOG44 in a 1:10 molar ratio using TransIT-X2 (Mirus Bio). After one day, cells were transferred to new dishes, and on the following day, hygromycin (100 μg/mL) was added to cells. Following selection, cells were maintained in 5 μg/mL of blasticidin and 100 μg/mL of hygromycin. For induction of mCherry or IFIT proteins, cells were treated with 500 ng/mL of doxycycline for 24 h.

MEFs from *Ifit2^-/-^*^11^ and Δ*Ifit3a/b* mice were prepared from day 13.5-14.5 embryos according to published protocols^54^. Isolated MEFs were maintained in DMEM supplemented with 10% heat-inactivated, fetal bovine serum (FBS) (Cytiva), 100 U/mL penicillin-streptomycin (Invitrogen), non-essential amino acids (Cellgro), and Glutamax (Gibco). Passage 0 (P0 MEFs were frozen or split 1:4 when approximately 80% confluent (around 3 days). To generate transformed MEFs, 5x10^6^ P1 primary MEFs were transfected with 10 μg of a plasmid (SV2)^55^ encoding the SV40 T antigen under control of a CMV promoter, using FuGene reagent (Promega) (3:1 μL FuGene to μg DNA ratio). Upon achieving confluence, MEF cultures were split 1:10. This process was repeated for approximately ten passages, at which time the transformed MEFs were frozen or used for experiments.

### IFIT siRNA knockdowns

Specific siRNAs against *ifit1* (Integrated DNA Technologies; 5’-UAGACGAACCCAAGGAGGCUCAAGCUU -3’), *ifit2* (Horizon Discovery, cat. # M-012582-01-0050), and *ifit3* (Integrated DNA Technologies, TriFECTa RNAi Kit - hs.Ri.IFIT3.13.1) were obtained from their respective manufacturers. A549 cells were seeded into 24-well plates. Twenty-four hours after seeding, cells were transfected with 20 pmol of siRNA in Lipofectamine 2000 Transfection Reagent (Invitrogen) and allowed to incubate for 24-hours before being used in subsequent infection experiments or being harvested for western blot to validate knockdown efficiency.

### Viral stocks and infections

VSV-GFP was a generous gift of Dr. John Rose^56^ (Yale University) and was propagated in BHK cells. For siRNA experiments, siRNA-treated A549 cells in 24-well plates were induced with 500 U/mL of IFN-α for 24 hours. Cells were then infected at a multiplicity of infection (MOI) of 0.01 for 16 h before harvesting, and virus was quantified by plaque assay. For ectopic overexpression experiments, Flp-In T-REx HEK293 cells in 24-well plates were induced with doxycycline for 24 h. Cells were then infected at an MOI of 0.01 for 16 h before harvesting, and virus was quantified by plaque assay. For high MOI experiments, Flp-In 293 cells in 96-well (imaging) or 24-well (western blotting) plates were infected at an MOI of 3.0, and cells were harvested or plates were imaged at 0, 2-, 4-, 6-, and 8-hours post-infection. For evaluation of structure guided mutations, 293T cells cultured in 24-well plates were transfected 6 h post-seeding with IFIT constructs. Eighteen hours post-transfection, cells were infected at an MOI of 0.05 for 16 h before imaging.

PIV3-GFP was obtained from Dr. Benhur Lee (Icahn School of Medicine at Mount Sinai)^57^. For siRNA experiments, siRNA-treated A549 cells in 24-well plates were induced with 500 U/mL of IFN-α for 24 h. Cells were then infected at an MOI of 0.01, and supernatant was harvested 40 hpi. Virus was quantified by TCID^50^ analysis. For ectopic expression experiments, Flp-In T-REx HEK293 cells in 24-well plates were induced with doxycycline for 24 h before being infected at an MOI of 0.1 for 48 h. Supernatant was then harvested, and virus was quantified by TCID^50^.

CVB3 stocks were generated by co-transfection of CVB3-Nancy infectious clone plasmids with a plasmid expressing T7 RNA polymerase (generous gifts of Dr. Julie Pfeiffer, University of Texas Southwestern Medical Center) as previously described^58^. For ectopic overexpression experiments, Flp-In T-REx HEK293 cells in 24-well plates were induced with doxycycline for 24 h before being infected at an MOI of 0.1. Supernatant was harvested 40 hpi and quantified by plaque assay.

### Western blotting and antibodies

Twenty-four hours post-transfection, cells were resuspended in supernatant and harvested, followed by centrifugation for 5 minutes at 2500 RPM. Cell pellets were washed with 1X PBS and lysed with 1X Bolt LDS Sample Buffer (Life Technologies, San Diego, CA) containing 5% β-mercaptoethanol at 98°C for 7 min. The lysed samples were centrifuged at 15,000 RPM for 2 minutes, followed by loading into a 4-12% Bolt Bis-Tris Plus Mini Protein Gels (Life Technologies, San Diego, CA) with 1X Bolt MOPS SDS Running Buffer (Life Technologies, San Diego, CA). Following electrophoresis, gels were wet transferred onto nitrocellulose membranes using a Mini Blot Module (Life Technologies, San Diego, CA). Membranes were blocked with PBS-T containing 5% bovine serum albumin (BSA) (Spectrum, New Brunswick, NJ), followed by incubation with primary antibodies (1:1000) for VSV-G [8G5F11] and VSV-N [10G4] (Kerafast), V5 [D3H8Q] (Cell Signaling Technology), or GAPDH [14C10] (Cell Signaling Technology). Membranes were rinsed three times in PBS-T and then incubated with the appropriate HRP-conjugated secondary antibodies (1:10000; Bio-Rad). Membranes were rinsed again three times in PBS-T and developed with SuperSignal West Pico PLUS Chemiluminescent Substrate (Thermo Fisher Scientific, Carlsbad, CA). Blots were imaged on a Bio-Rad ChemiDoc MP using the Image Lab Software suite (Bio-Rad).

### Plasmids, constructs, and molecular cloning

The coding sequences of mouse IFIT2 (NCBI accession BC050835) and mouse IFIT3 (NCBI accession BC089563) were cloned separately into the pcDNA5/FRT/TO backbone (Invitrogen, Carlsbad, CA) with an N-terminal 3xFLAG tag or the pQCXIP backbone (Takara Bio, Mountain View, CA) with an N-terminal HA tag, respectively, and both were cloned into the pcDNA5/FRT/TO backbone (an N-terminal HA tag, followed by IFIT3, a p2a site, a 3xFLAG tag, and IFIT2). IFIT2 and IFIT3 point mutants were generated using overlapping stitch PCR and cloned into their respective backbones. The 5’ UTR and coding sequence for each VSV gene (GenBank accession number NC_038236.1) was cloned into the pQCXIP backbone with a C-terminal V5 tag. Following 5’ RACE, “short” 5’ UTR constructs were generated by truncating the pQCXIP 5’ UTR sequence. The “long” and “short” backbone constructs expressing VSV-P-V5 were used to further clone the VSV-P fluorescence reporter plasmids by subcloning GFP in between the VSV-P 5’ UTR and V5 tag. The mCherry normalization construct was generated by subcloning mCherry and a C-terminal 3xFLAG tag into the pcDNA5/FRT/TO backbone. All additional reporter constructs (including VSV-P truncations, UTR length constructs, and viral 5’ UTRs [RABV (NCBI accession NC_001542)]; [PIV3 (NCBI accession NC_001796)]; [CVB3 (NCBI accession NC_038307)]) were cloned using primers and inserted into the “short” UTR reporter backbone upstream of GFP-V5. All generated plasmids were sequenced across the entire inserted region to verify that no mutations were introduced during the cloning process. Plasmids and primers used in this study can be found in **Supplementary Table 3**. Gene Fragments were ordered from Twist Bioscience (South San Francisco, CA). All newly created plasmids will be made available upon request.

### Protein expression and purification

For expression of the IFIT2-IFIT3 complex, we cloned codon-optimized genes encoding *M. musculus Ifit2 and Ifit3* into separate plasmid vectors for expression in *E. coli*, with *Ifit2* cloned into UC Berkeley Macrolab vector 2-BT (Addgene #29666; ampicillin resistant) encoding a TEV protease-cleavable His6-tag, and *Ifit3* cloned into UC Berkeley Macrolab vector 13S-A (Addgene #48323; spectinomycin resistant) with no tag.

Plasmids were co-transformed into *E. coli* Rosetta pLysS cells (EMD Millipore) and grown overnight at 37°C in LB plus carbenicillin and spectinomycin. Saturated overnight cultures were used to inoculate six 1 L cultures of 2XYT media plus carbenicillin and spectinomycin, and cultures were grown at 37°C with shaking at 180 RPM to an OD600 of 0.8. Protein expression was induced by the addition of 0.25 mM IPTG, then cultures were shifted to 20°C and grown another 16 h with shaking. Cells were harvested by centrifugation and resuspended in Nickel Wash Buffer (20 mM Tris-HCl pH 7.5, 500 mM NaCl, 20 mM imidazole pH 8.0, 2 mM beta-mercaptoethanol, and 10% glycerol).

For protein purification, resuspended cells were lysed by sonication (Branson Sonifier), then cell debris was removed by centrifugation at 14,000 RPM in a JA-17 rotor in an Avanti J-E centrifuge (Beckman Coulter) for 30 min. Clarified lysate was passed over a nickel column (5 mL HisTrap HP, Cytiva) in Nickel Wash Buffer, then bound protein was eluted with Nickel Elution Buffer (20 mM Tris-HCl pH 7.5, 75 mM NaCl, 250 mM imidazole pH 8.0, 2 mM beta-mercaptoethanol, and 10% glycerol). Eluted protein was concentrated and buffer-exchanged into Nickel Elution Buffer containing 20 mM imidazole (Amicon Ultra, EMD Millipore), and the His^6^-tag on IFIT2 was cleaved by addition of 1:10 w/w ratio of purified TEV protease (S219V mutant, purified in-house from expression vector pRK793; AddGene #8827)^59^, followed by incubation at 4°C for 48 h. The reaction mixture was passed over a nickel column to remove cleaved His^6^-tags, uncleaved IFIT2, and His^6^-tagged TEV protease. The flow-through was concentrated, then passed over a size exclusion column (Superdex 200 Increase, Cytiva) in Size Exclusion Buffer (20 mM Tris-HCl pH 7.5, 150 mM NaCl, and 1 mM DTT) and fractions containing both proteins were pooled and concentrated.

### Cryo-EM grid preparation

Prior to use, UltrAuFoil 1.2/1.3 300 mesh grids were plasma cleaned for 12 sec using a Solarus II plasma cleaner (Gatan). Purified IFIT2-IFIT3 at 3 mg/mL was applied to the grid in a 3 μL drop within the environmental chamber adjusted to 4°C temperature and approximately 95% humidity in a Vitrobot Mark IV (ThermoFisher Scientific). After a 4 sec incubation, the grids were blotted with a blot force of 4 for 4 sec; the sample was then plunged frozen into liquid nitrogen-cooled liquid ethane.

### Cryo-EM data acquisition and image processing

All data was acquired at the UCSD Cryo-EM Facility on a Titan Krios G3 electron microscope (Thermo Fisher Scientific) operating at 300 kV and equipped with a Gatan BioContinuum energy filter. All images were collected at a nominal magnification of 165,000x in EF-TEM mode (with a calibrated pixel size of 0.854 Å) on a Gatan K2 detector using a 20-eV slit width and a cumulative electron exposure of ∼65 electrons/Å^2^ over 50 frames **(Supplementary Table 1)**. Data were collected automatically using EPU (Thermo Fisher Scientific) with aberration-free image shift with a defocus range of -1 to -2.5 µm. Data collection was monitored live using cryoSPARC Live (Structura Bio)^60^ where movies were patch motion-corrected and patch CTF-estimated on the fly. Micrographs with a CTF estimation worse than 7 Å and/or a cumulative motion of more than 150 pixels were discarded.

An initial 484,841 particle picks were obtained using crYOLO^61^ picker using a general model, and these picks were imported into cryoSPARC^60^. Particles were extracted with a box size of 384 pixels and Fourier-cropped to 96 pixels at (3.34 Å/pixel). These particles were subjected to three rounds of two-dimensional (2D) classification, where classes with proteinaceous features were chosen to move forward. The selected particles were subjected to an *ab initio* reconstruction to generate a starting model and carried forward to a non-uniform (NU) refinement using C1 symmetry. These particles were then re-extracted at a box size of 384 pixels with a Fourier crop to 128 pixels (1.708 Å/pixel) and a second NU refinement was performed. The particle stack was then subjected two rounds of a two-class heterogeneous refinement with one volume being the volume from the previous NU refinement, and the other volume from EMD-4877 (20S proteasome)^62^, followed by NU refinement. In each round, the particles that contributed to the best volume that resembled the IFIT2-IFIT3 dimer were selected. A final 3-class heterogenous refinement was performed using two IFIT2-IFIT3 volumes and a 20S proteasome volume. The particles associated with the volume that showed the best secondary structure features was selected, and NU refinement was performed, resulting in a 3.51 Å resolution map. These particles were then re-extracted at a box size of 384 pixels with no Fourier cropping (0.854Å/pixel). These particles were then NU refined, resulting in a 3.57 Å resolution map. Due to heterogeneity that was observed in the map, particles were then exported from cryoSPARC into RELION^63^, where they were extracted at a box size of 256 with a Fourier crop of 64 (3.34Å/pixel). These particles were subjected to a round of 2D classification in which obvious junk classes were discarded. Selected particles were then re-extracted at a box size of 384 (0.854Å/pixel) and subjected to 3D auto refinement, CTF refinement, and a second 3D auto refinement^64^. The particles were then Bayesian particle polished^65^ before another round of two 3D auto refinements and a CTF refinement. This final particle stack was then imported back into cryoSPARC for a final NU refinement that resulted in a 3.22 Å resolution map **(Supplementary Table 1)**.

To generate a starting model, we used ModelAngelo^66^ and supplied our final 3.22Å map and sequence files for IFIT2 and IFIT3. This resulting model was then iteratively real-space refined using Phenix^67^ and manually adjusted in COOT^68^. After the final refinement, the model was checked for accuracy in COOT **(Supplementary Table 1)**.

### Evolutionary analyses

For evolutionary analyses of primate and rodent IFIT2 and IFIT3, Uniprot reference protein sequences for human IFIT2, human IFIT3, mouse IFIT2, and mouse IFIT3 were used as a search query against NCBI’s non-redundant (NR) database using tBLASTn^69^. Searches were restricted to simian primates and the Muroidea superfamily of rodents respectively. For each species, the nt sequence with the highest bit score was downloaded and aligned to the human or mouse ORF nt sequence using MAFFT^70^ implemented in Geneious software (Dotmatics; geneious.com). Poorly aligning sequences or regions were removed from subsequent analyses. Accession numbers of final sequences used for analyses are in **Supplementary Table 2**. Using these aligned sequences, FUBAR^71^ was performed on Datamonkey.org using 50 grid points and a 0.5 concentration parameter of the Dirichlet prior to infer codons evolving under positive and negative selection. Codons with a posterior probability of 0.9 or higher are in **Supplementary Table 2**. PAML^72^ was used to infer gene-wide positive selection, as well as codon-based estimates of positive selection. Aligned sequences were analyzed using the NS sites models disallowing (M7) or allowing (M8) positive selection. The p-value reported is the result of a chi-squared test on twice the difference of the log likelihood (lnL) values between the two models using two degrees of freedom. Analyses were performed using two models of frequency (F61 and F3x4) and both sets of values are reported in **Supplementary Table 2**. For each codon model, we confirmed convergence of lnL values by performing each analysis using two starting omega (dN/dS) values (0.4 and 1.5). Positively selected codons with a posterior probability greater than 0.90 using a Bayes Empirical Bayes (BEB) analysis and the F61 codon frequency model are in **Supplementary Table 2**.

### eCLIP experimental methods

Flp-In T-REx HEK293 cells in 10 cm culture dishes were induced with doxycycline for 24 h. Cells were then infected at an MOI of 3.0, and dishes were crosslinked using a UV cross-linker (254 nm; CL-1000 from UVP/Analytik Jena) at 400 mJ/cm^2^. Downstream sample processing and eCLIP were performed as previously described^73^, using antibodies against FLAG (M2 / F1804, Sigma) and HA (HA.11 / 901502, Biolegend) for 293T experiments, and IFIT2 (PA3-845, ThermoFisher) and IFIT3 (ABF1048, Millipore) for MEF experiments. Most experiments were performed in biological duplicate, excepting the uninfected 293T samples (which were single replicates).

### eCLIP computational analysis

Standard processing of eCLIP data was performed as previously described^73^, with mapping performed to a custom genome index that included both the VSV genome and either hg19 (for 293T experiments) or mm10 (for MEF experiments). Data (**Fig. 3**, **Extended Data** Fig. 5) are plotted as normalized reads per million (RPM), where reads per million are normalized to density of reads mapped to viral and human genomes.

### Animal Ethics statement and generation of *Ifit3a/b* DKO mice

Experiments was carried out in accordance with the recommendations in the Guide for the Care and Use of Laboratory Animals of the National Institutes of Health. The protocols were approved by the Institutional Animal Care and Use Committee at the Washington University School of Medicine (Assurance number A3381-01). Wild-type C57BL/6J were commercially obtained from Jackson Laboratories (Strain #000664; Bar Harbor, ME). To generate *Ifit3* deficient mice, single guide RNAs (sgRNAs) were designed to target exon two in *Ifit3a* and *Ifit3b*. Two sgRNAs were chosen that target conserved sequences between *Ifit3a* and *Ifit3b*: sgRNA-4, 5’-ATTTCACCTGGAATTTATTCNGG-3’, and sgRNA-30, 5’-AATGGCACTTCAGCTGTGGANGG -3’. Two additional sgRNAs were chosen that target only *Ifit3a* due to polymorphisms: sg-RNA-21, 5’-AATTCGTCGACTGGTCACCTNGG -3’, and sgRNA-22, 5’-ATTCGTCGACTGGTCACCTGNGG -3’. The sgRNAs were selected based on their low off-target profile. Guide RNAs and Cas9 protein were complexed and electroporated concurrently into C57BL/6J zygotes. Using this approach, we identified a mouse that had both *Ifit3a* and *Ifit3b* targeted (22 nucleotide (nt) and 20 nt frameshift deletions, respectively), two mice in which only *Ifit3a* was targeted (2 nt and 119 nt frameshift deletions), and two mice in which only *Ifit3b* was targeted (1 nt and 2 nt frameshift insertions). After genotyping and two rounds of backcrossing, five founder lines (*Ifit3a* del22/*Ifit3b* del20; *Ifit3a* del2; *Ifit3a* del119; *Ifit3b* ins1; and *Ifit3b* ins2) were generated. The generation of gene-edited mice was accomplished with the aid of the Genome Engineering and iPSC center and Department of Pathology Micro-Injection Core (Washington University School of Medicine).

### 5’ RACE (Rapid Amplification of cDNA Ends)

HEK293T cells were maintained as described above and subcultured into 6-well plates (Genesee, El Cajon, CA) in 2 mL of complete media 24 h before transfection. Cells were transiently transfected with 2500 ng of total DNA and 7.5 μL of TransIT-X2 (Mirus Bio, Madison, WI) following the manufacturer’s protocol. 24 hours post-transfection, cells were harvested and pelleted; cell pellets were washed with 1x PBS, pelleted again, and supernatant was aspirated. RNA was extracted from pellets using the Takara Bio NucleoSpin RNA Plus kit following the manufacturer’s protocol. Downstream processing of RNA was performed using the Takara Bio SMARTer RACE 5’/3’ Kit according to the manufacturer’s protocol.

### Fluorescent reporter assay

Cells (HEK293T or inducible Flp-In lines) were maintained as described above and subcultured into 24-well plates for transfection. Transfections were performed as described above with the exception of the addition of 100 ng of an mCherry-expressing DNA plasmid, resulting in 600 ng of total DNA transfected along with 1.8 μL of TransIT-X2. 24 hours post-transfection, cells were imaged using the BioTek Cytation 5 Cell Imaging Multimode Reader. Four images were taken at fixed positions in each well using a 20X objective lens, with each condition in two replicates; both GFP and RFP images were collected for each well position. Non-transfected wells were imaged for use as background subtraction. Images were pre-processed in the Gen5 3.1 software package (BioTek) using the default rolling ball algorithm settings, and mean fluorescence values for GFP and RFP were quantified by the Gen5 software.

### Image analyses

Normalized GFP intensity for each image was calculated as follows: (experimental GFP signal – background GFP signal) / (experimental RFP signal – background RFP signal). Background GFP and RFP signals are the average of the quantified pre-processed values from 8 total images taken of non-transfected wells. Once the normalized GFP intensity was calculated for each image, we averaged the normalized GFP intensity of each set of four images (*i.e*. for each well). Finally, we calculated the relative GFP intensity for each well by dividing the average normalized GFP intensity from each well by the average of the two IFIT-untreated wells, thus representing each data point as relative to 100%.

### qRT-PCR

HEK293T cells were maintained as described above in 24-well plates. Cells were harvested in 1x PBS and pelleted at 500 x g for 1 minute. RNA was then extracted using the New England BioLabs Monarch Total RNA Miniprep Kit according to the manufacturer’s protocol. 100 ng of RNA was then subjected to reverse transcription using the Applied Biosystems High-Capacity cDNA Reverse Transcription Kit according to the manufacturer’s protocol. The resulting cDNA was then used to conduct quantitative PCR on the Applied Biosystems StepOnePlus machine with gene-specific primers (GFP-F, 5’-CCGACCACTACCAGCAGAACAC -3’; GFP-R, 5’-GGACCATGTGATCGCGCTTCTC -3’; 18S-F, 5’-TCGCTCGCTCCTCTCCTACTTG -3’; 18S-R, 5’-GCTGACCGGGTTGGTTTTGATCTG -3’) and the Luna Universal qPCR Master Mix according to the manufacturer’s protocol.

### Statistical analyses

Statistical analyses and data visualization were performed using GraphPad Prism 10 software. Tests were performed as indicated in figure legends. All error bars represent SEM.

### Material availability statement

All unique materials/reagents generated in this study are available through the lead contact upon request.

## DATA AVAILABILITY

All data reported in this paper will be shared by the lead contact upon request. The cryo-EM structure has been deposited under PDB 9MK9 and EMDB code EMD-48323. Sequencing data from eCLIP experiment have been deposited in GEO record GSE284636.

## ACKNOWLEDGEMENTS

We thank Laura A. VanBlargan for her contribution to the initial design and testing of *Ifit3*-deficient mice and cells. We also thank Drs. Eric Bennett, Andy Mehle, Benhur Lee, and Sara Cherry, as well as members of the Daugherty and Biering Labs, for helpful discussions and Drs. Patrick Mitchell and Scott Biering for comments on the manuscript.

The authors acknowledge support from the National Institutes of Health (M.D.D.: K22 AI119017, R35 GM133633; E.L.V.N.: R00 HG009530; G.W.Y.: U24 HG009889, RF1 MH126719, R01 HG011864, R01 HG004659, subaward from U24 HG011735; K.D.C.: R35 GM144121; S.K.: T32 GM133351), the Burroughs Wellcome Fund Investigators in the Pathogenesis of Infectious Disease Program (to M.D.D.).

## AUTHOR INFORMATION

### Contributions

Conceptualization, D.R.G., M.D.D.; Methodology, D.R.G., C.T., A.D., E.E., J.D.W., M.D.D.; Investigation, D.R.G., C.T., B.T., S.K., A.D., E.E., J.D.W., P.F., J.P., K.H., B.A.Y., K.M.E., Q.L., S.R.H., J.B.C., E.L.V.N.; Formal Analysis, D.R.G., M.D.B., K.D.C., E.L.V.N., M.D.D.; Resources, J.B.C., K.D.C., M.S.D.; Writing – Original Draft, D.R.G., B.C., K.D.C., M.D.D.; Writing – Review & Editing, All authors except S.R.H., who has tragically passed away since making important initial discoveries for this work; Visualization, D.R.G., B.C., M.D.B., K.D.C., E.L.V.N., M.D.D.; Supervision, K.D.C., M.S.D., G.W.Y., M.A.H., E.L.V.N., M.D.D.; Funding Acquisition, M.D.D., E.L.V.N., G.W.Y., K.D.C.

### Corresponding authors

Requests for further information and resources should be directed to and will be fulfilled by the lead contact, Matthew D. Daugherty.

## EXTENDED DATA

**Extended Data Fig. 1:**
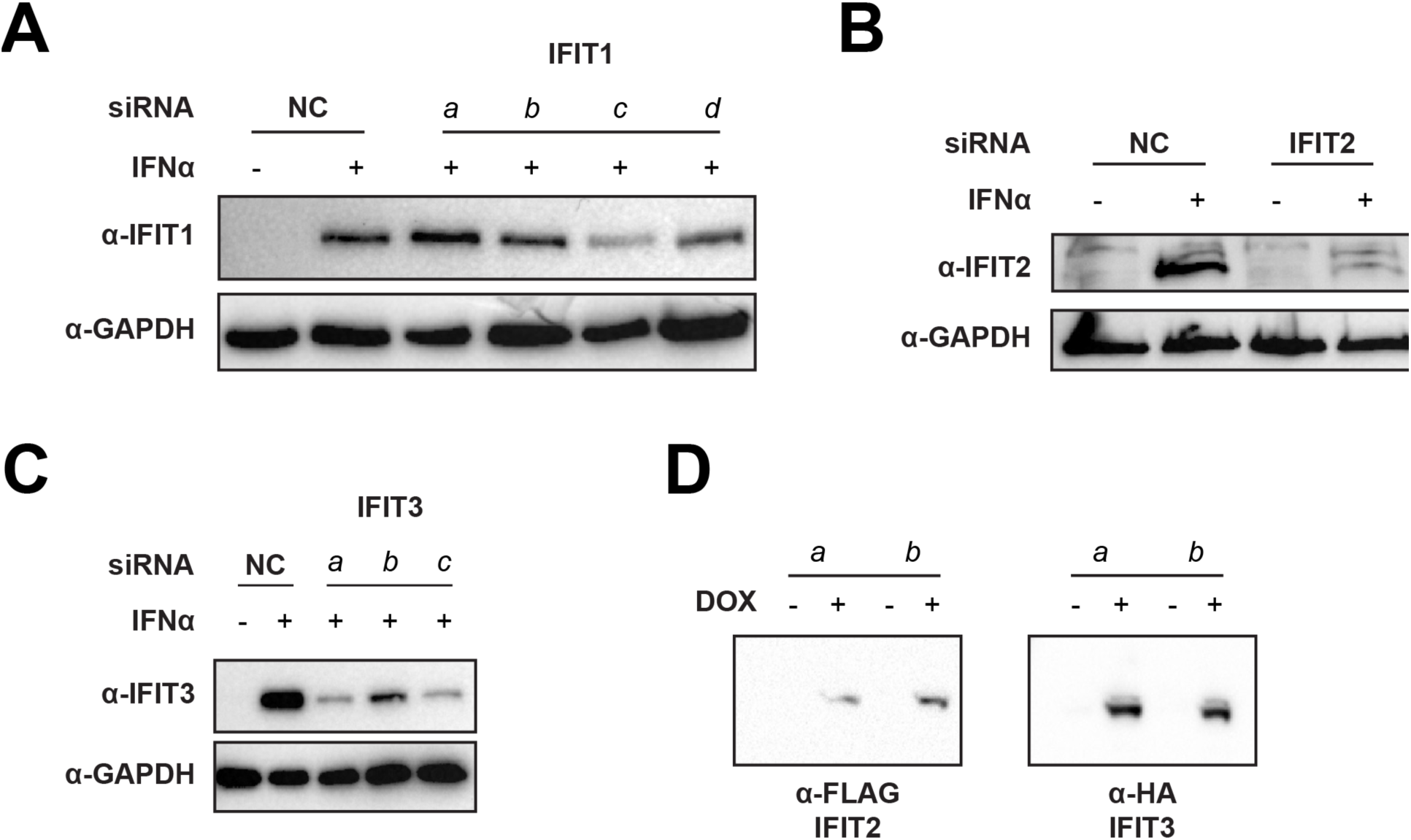
siRNA knockdown of IFITs in A549 cells and expression of IFIT2 and IFIT3 in FlpIn lines, related to. Fig. 1. (**A-C**) A549 cells were treated with a non-targeting siRNA as a negative control or siRNA targeting human (**A**) *IFIT1*, (**B**) *IFIT2*, or (**C**) *IFIT3*. Twenty-four hours post-treatment, cells were harvested and subsequently processed and analyzed by western blotting. Representative images shown; 2 independent experiments. For downstream infection experiments, siRNA IFIT1-c and siRNA IFIT3-a were used. (**D**) We generated inducible Flp-In T-REx HEK293 cell lines co-expressing mouse IFIT2 and IFIT3. Cells were induced with 500 ng/mL DOX for 24 hours before harvesting cell pellets for analysis by western blot. For all associated downstream experiments, Flp-In line *b* was used.

**Extended Data Fig. 2:**
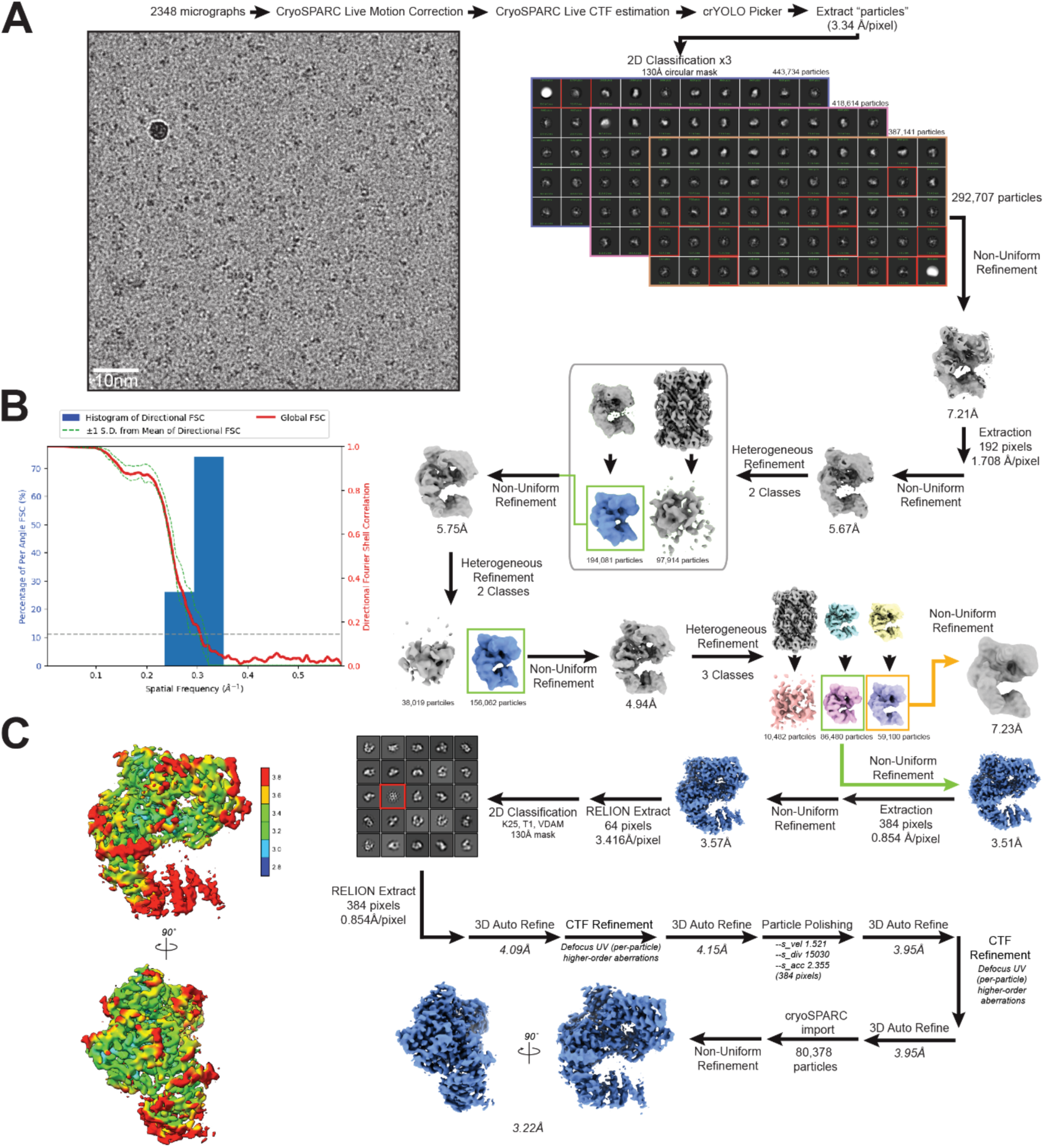
Cryo-EM data processing for IFIT2-IFIT3, related to Fig. 2. (**A**) Workflow for structure determination. See **Methods** for a detailed explanation of data processing. For 2D classes, classes outlined in red were excluded from later steps. For 3D classes, classes outlined in green were selected for later steps. The final refined map showed a resolution of 3.22 Å (gold-standard FSC criterion of 0.143). (**B**) Directional Fourier Shell Correlation analysis for the final map, calculated by 3DFSC^74^. (**C**) Two views of the final map, colored by local resolution (blue: 2.8 Å or better; red: 3.8 Å or worse).

**Extended Data Fig. 3:**
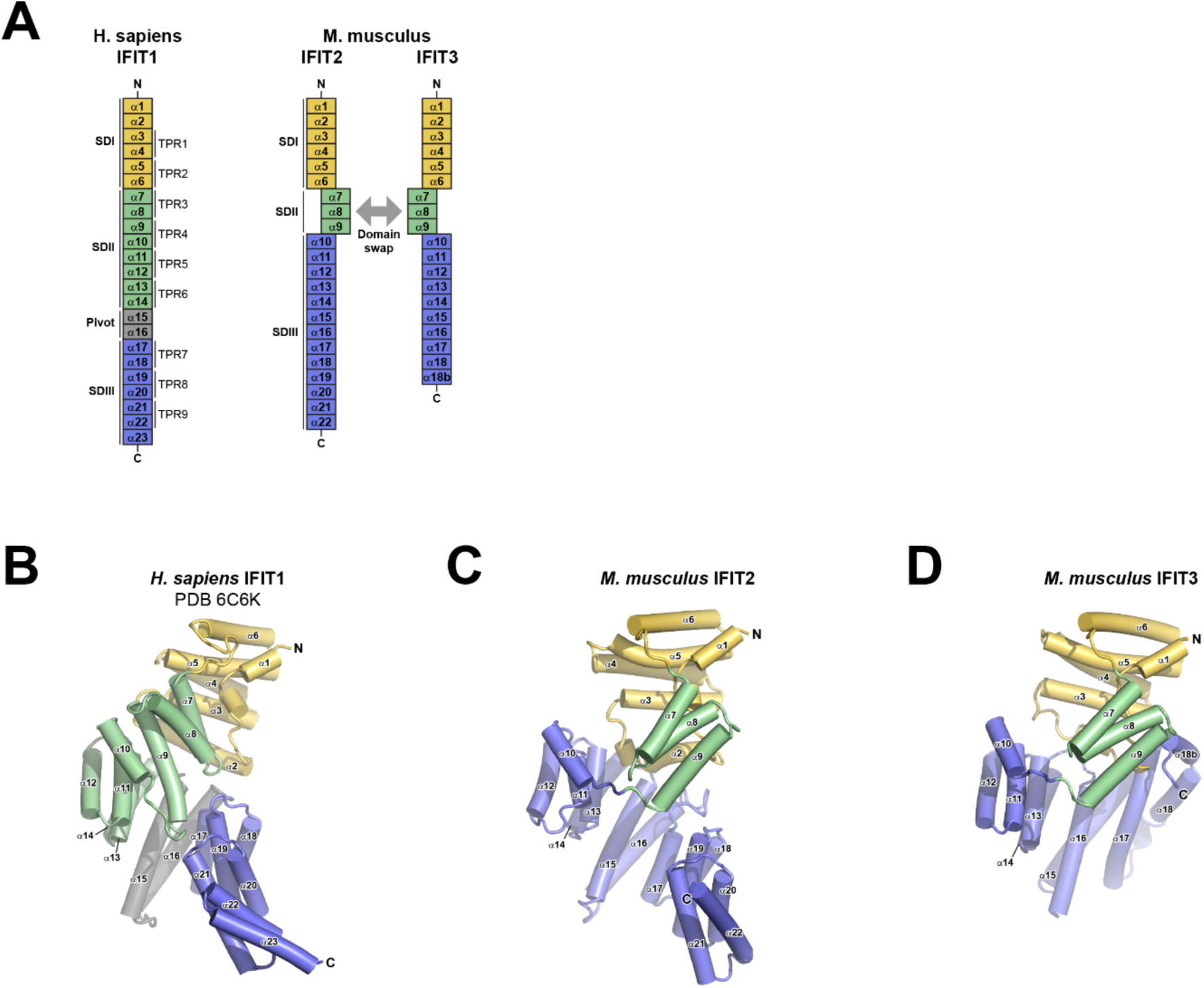
Comparison of IFIT structures, related to. Fig. 2. (**A**) Representation of human IFIT1 and mouse IFIT2 and IFIT3 domains. (**B-D**) Representations of (**B**) human IFIT1, (**C**) mouse IFIT2, and (**D**) mouse IFIT3. (**C-D**) The white, semitransparent helices represent the domain-swapped helices during IFIT2-IFIT3 heterodimer formation.

**Extended Data Fig. 4:**
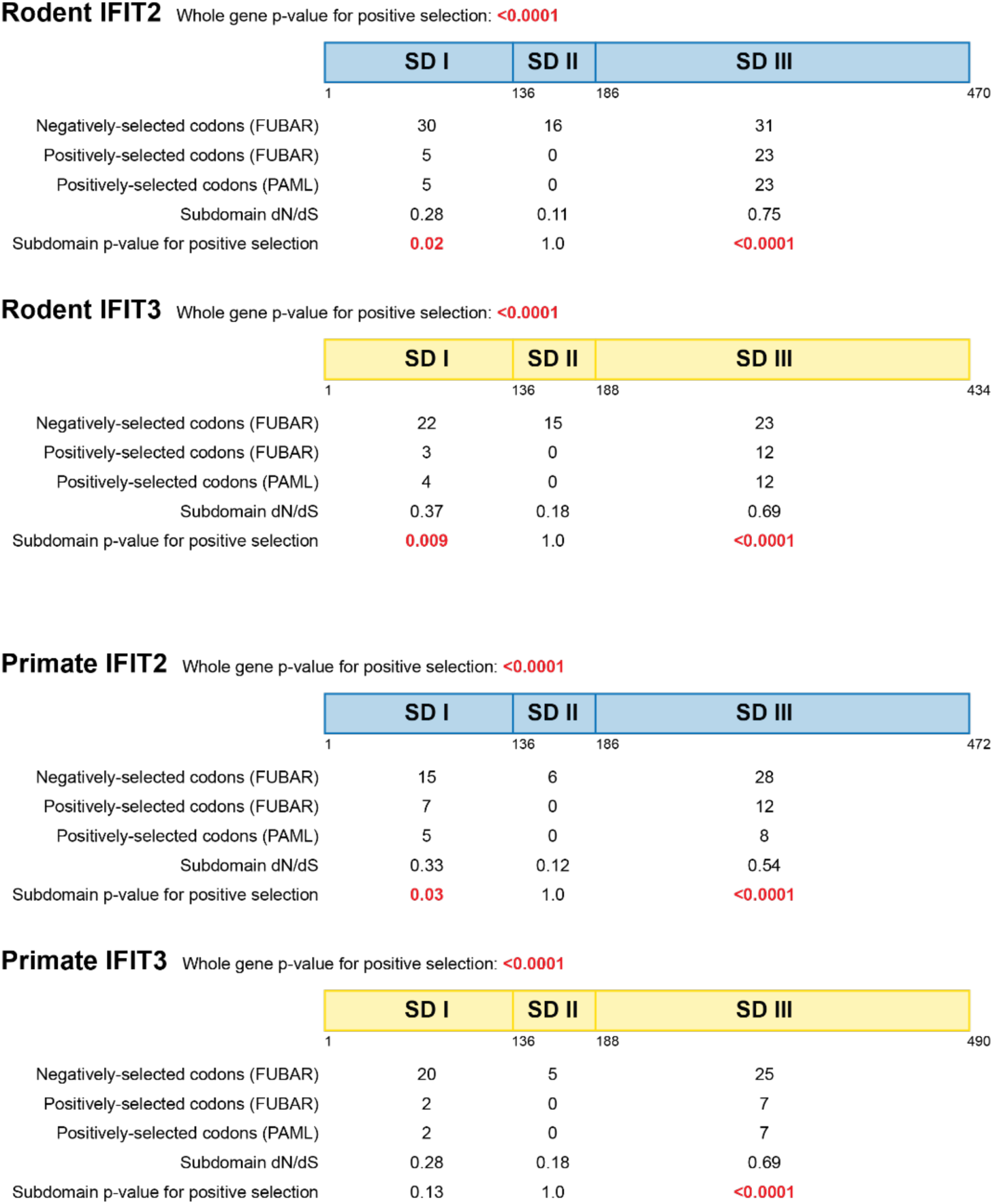
Evolutionary analysis, related to. Fig. 2. Whole gene and subdomain evolutionary analyses on rodent and primate IFIT2 and IFIT3. Domain boundaries for each subdomain (SD) are indicated by amino acid numbers below each schematic. Statistical evidence for positive selection, determined by PAML software, is shown for each gene and each subdomain, with statistically significant p-values shown in bold red text. Overall dN/dS (omega) values for each subdomain are also shown. The number of specific codons within each subdomain that show statistical evidence for positive and negative selection, as determined by PAML and FUBAR software as indicated. Additional statistics and analysis parameters, complete lists of codons, and accession numbers and species of sequences analyzed are found in Table S2.

**Extended Data Fig. 5:**
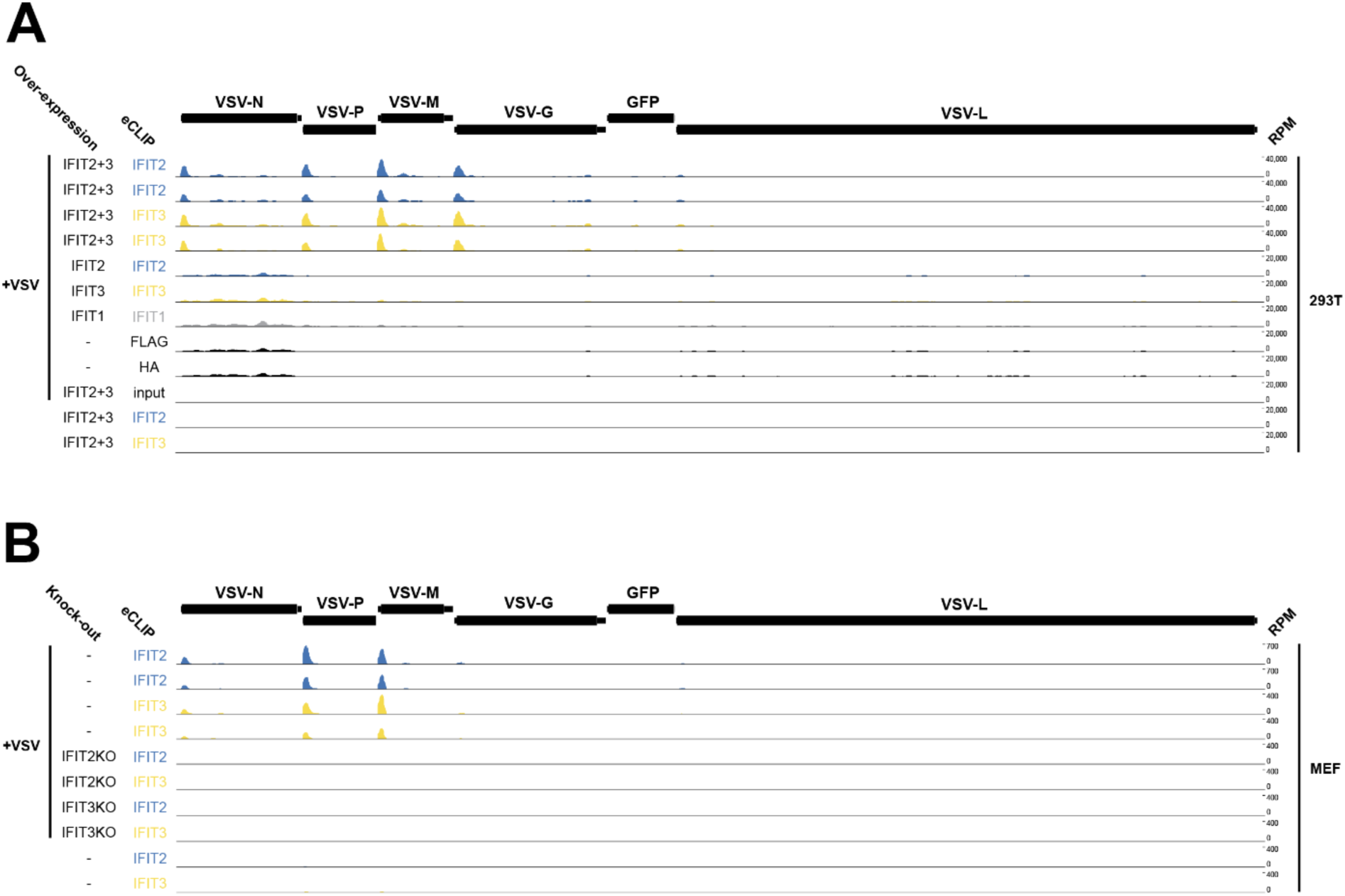
Complete set of eCLIP data, related to. Fig. 3. Full VSV genome coverage for additional samples from the eCLIP experiment described in Fig. 3, including uninfected controls, for both (**A**) human IFIT-expressing Flp-In T-REx HEK293 lines and (**B**) mouse wild-type and *Ifit2* or *Ifit3* knockout MEFs. Values shown are normalized reads per million (RPM) out of all reads mapped to the VSV genome.

**Extended Data Fig. 6:**
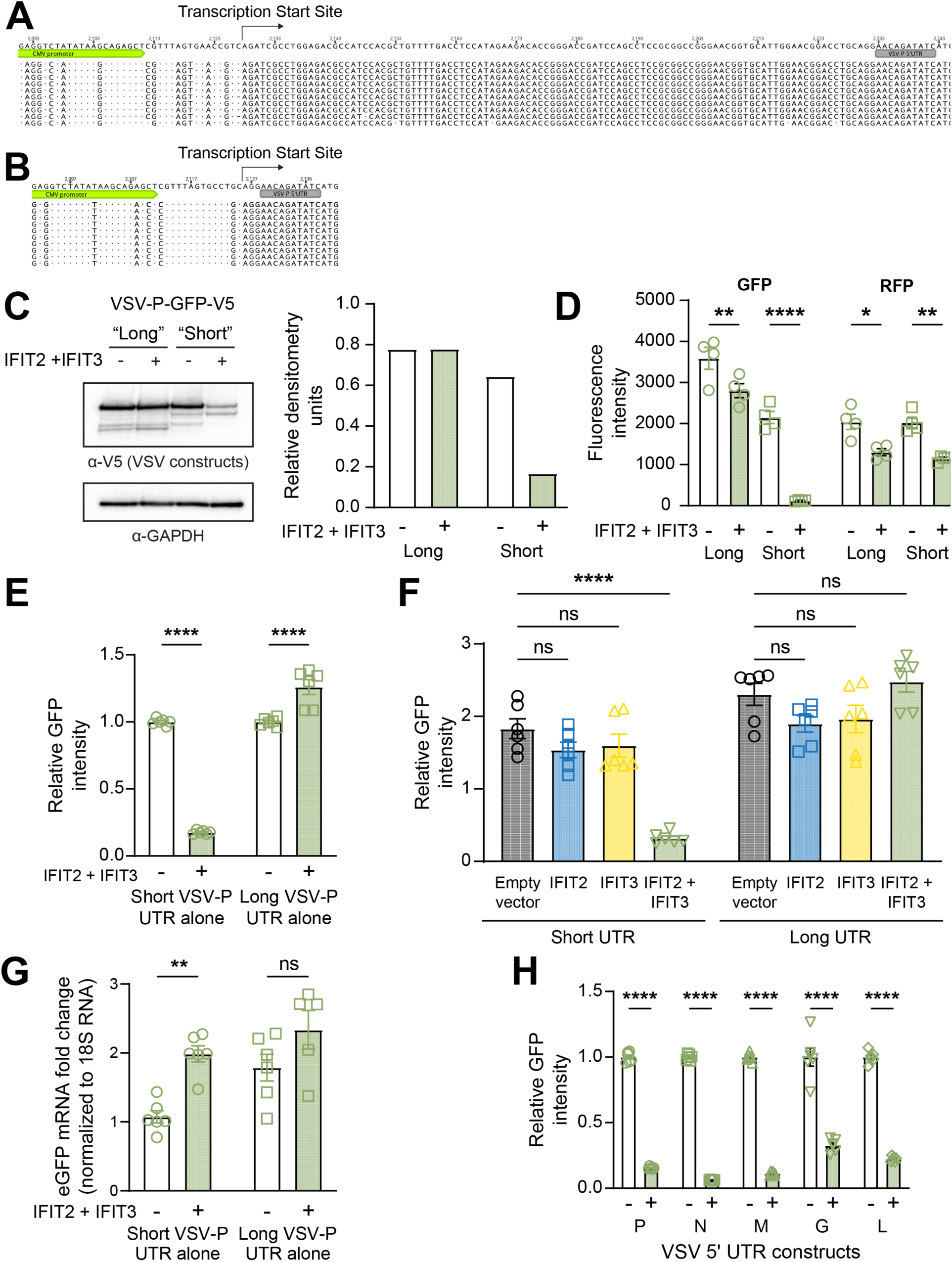
IFIT2-IFIT3 recognizes short viral 5’ UTRs, related to. Fig. 4. (**A-B**) Plasmids expressing either a (**A**) long plasmid-derived 5’ UTR or (**B**) no plasmid-derived 5’ UTR immediately upstream of the VSV-P 5’ UTR were transfected into 293T cells. Twenty-four hours post-transfection, cells were harvested, and RNA was isolated for 5’ RACE experiments. Ten clones from each construct were sequenced, and the transcription start site is marked for each plasmid. (**C**) Experiment was carried out as described in Figure 4C, but instead of imaging, cells were harvested and lysates were analyzed by western blotting. Densitometry values were calculated using ImageJ. **(D)** Raw values from normalized data in panel C. (**E-F**) 293T cells were transfected with plasmids expressing the 10 nt VSV-P 5’ UTR and eGFP with either a short or long plasmid-derived 5’ UTR upstream of the viral sequence, co-transfected in the absence or presence of (**E**) IFIT2-IFIT3 or (**F**) IFIT2 or IFIT3 alone. Experiments were performed as described in Figure 4F-G. (**G**) Cells were transfected as in panel E, and 24 hours post-transfection, cells were harvested, RNA was extracted, and qRT-PCR was conducted. Data are pooled from two independent experiments, and data are normalized to the short VSV-P UTR alone, no IFIT2-IFIT3 condition. (**G**) Cells were transfected with plasmids expressing GFP and the 5’’ UTR of each VSV gene in the absence or presence of IFIT2-IFIT3. Experiments were performed as described in Figure 4F-G. Statistical analyses: Data are represented as mean ± SEM. (**D, E, G, H**) Ordinary two-way ANOVA with Šídák’s multiple comparisons test and a single pooled variance. * = p<0.05, ** = p<0.01, **** = p<0.0001, ns = not significant; (**F**) ordinary two-way ANOVA with Dunnett’s post-test and a single pooled variance. Comparisons are to the empty vector condition within each column. **** = p<0.0001, ns = not significant.

## SUPPLEMENTARY INFORMATION

Supplementary Table 1. Cryo-electron microscopy data collection and structure determination, related to Figure 2

**Supplementary Table 2. Excel file containing evolutionary analyses-related information, related to Figure 2 and S4.** Individual tables contain accession numbers and species for sequences analyzed, complete lists of codons determined to be evolving under purifying and positive selection, and parameters and statistics for whole-gene and subdomain analyses.

**Supplementary Table 3. Excel file containing information on plasmids and primers, related to Experimental Procedures.** Spreadsheet containing all plasmids used in this study, along with a brief description and primers and/or Gene Fragments used for cloning.

