## Supplementary Table 1 for "Short 5’ UTRs serve as a marker for viral mRNA translation inhibition by the IFIT2-IFIT3 antiviral complex"

**Supplementary Table 1. Cryo-electron microscopy data collection and structure determination, related to Fig. 2**

| | |
|---|---|
| **Data Collection** | |
| Magnification | 165,000 |
| Voltage (kV) | 300 |
| Movies | 2355 |
| Spherical Aberration (mm) | 2.7 |
| Electron Exposure ($e^-$/Å$^2$) | 65 |
| Defocus range (μm) | -1 to -2.5 |
| Pixel size (Å, Physical/Digital) | 0.854 |
| Energy Filter Slit Width (eV) | 10 |
| **Map Statistics and Post-Processing** | |
| Symmetry imposed | C1 |
| Map Resolution (Å) | 3.22 |
| Local resolution range for 75% of voxels (Å$^2$) | 6.207 |
| Local resolution range (Å$^2$) | 2.9 - 26.6 |
| Map sharpening B factor (Å$^2$) | 107.7 |
| Map sharpening method | B-Factor |
| Q-Score | 0.58 |
| **Model Statistics and Validation** | |
| Model composition | |
| Non-hydrogen atoms | 6798 |
| Protein residues | 822 |
| Water | 47 |
| R.M.S deviations | |
| Bond length (Å) | 0.003 |
| Bond angles (°) | 0.592 |
| MolProbity score | 1.37 |
| MolProbity Clashscore | 6.62 |
| CaBLAM (% outliers) | 1.46 |
| Rotamer outliers (%) | 0.00 |
| Ramachandran Plot | |
| Favored | 97.67 |
| Allowed | 2.33 |
| Outliers | 0.0 |
